# scMusketeers: Addressing imbalanced cell type annotation and batch effect reduction with a modular autoencoder

**DOI:** 10.1101/2024.12.15.628538

**Authors:** Antoine Collin, Simon J. Pelletier, Morgane Fierville, Arnaud Droit, Frédéric Precioso, Christophe Bécavin, Pascal Barbry

## Abstract

The growing number of single-cell gene expression atlases available offers a conceptual framework for improving our understanding of physio-pathological processes. To take full advantage of this revolution, data integration and cell annotation strategies need to be improved, in particular to better detect rare cell types and by better controlling batch effects in experiments. scMusketeers is a deep learning model that optimises the representation of latent data and solves both challenges. scMusketeers features three modules: (1) an autoencoder for noise and dimensionality reductions; (2) a focal loss classifier to enhance rare cell type predictions; and (3) an adversarial domain adaptation (DANN) module for batch effect correction. Benchmarking against state-of-the-art tools, including the UCE foundation model, showed that scMusketeers performs on par or better, particularly in identifying rare cell types. It also allows to transfer cell labels from single-cell RNA sequencing to spatial transcriptomics. With its modular and adaptable design, scMusketeers offers a versatile framework that can be generalized to other large-scale biological projects requiring deep learning approaches, establishing itself as a valuable tool for single-cell data integration and analysis.

## 1 Introduction

Single-cell gene expression atlases are now essential in many biological studies about physio-pathological processes, cell differentiation, organ development, and the impact of mutations in diseases. Two key tasks in single-cell atlas construction are integration [1] and cell annotation [2]. Deep learning has recently advanced both tasks, often treated independently. However, both face similar challenges inherent to singlecell data: high volume, large dimensionality, technical and biological noise, complex expression patterns, and imbalanced settings.

In a single-cell atlas each gene is considered as a feature, resulting in a highdimensional expression space with inflated distances and sparse data distribution [3]. Dimensionality reduction is thus crucial to uncover structure in the data and cell similarities. Additionally, technical noise varying across experiments can obscure biological signals. Integration methods address these issues by creating a latent space of reduced dimension that captures core biological information while minimizing batch effects. Strictly speaking, this so-called “integration”, rather corresponds to an harmonization that deals with varying batch effect strengths across datasets (e.g., different labs, technologies, or cell quantification methods) [4].

Embedding space reconstruction during integration primarily follows two approaches. The first encourages similar cells from different datasets to cluster together. This is well illustrated by Harmony [5], a widely used method available in the Seurat and Scanpy workflows. It iteratively corrects principal components to group similar cells while maximizing batch diversity and minimizing technical variations. The second approach creates a compressed latent space capable of data recovery with batch identity regressed out. This method has advanced significantly with deep learning. For instance, Universal Cell Embedding (UCE) [6] leverages foundation models, training a transformer on a large cell atlas corpus across species using protein-level gene representations. This produces a unified biological latent space applicable to any cell, tissue, or species.

The purpose of cell type annotation is to identify biologically relevant cell groups. It is primarily a semi-automatic process, using unsupervised clustering and marker gene identification, which are validated by experts [7]. Various methods aim to fully automate this process, including marker-based, similarity-based, and machine learning approaches [2]. Marker-based models use external databases of marker genes for prior modeling or to compute cell type scores [8, 9]. Similarity-based methods compute similarity scores between annotated and unannotated cells, such as scmap, which uses correlation and cosine similarity to propagate labels [10]. Machine learning methods, like Celltypist, train classifiers on annotated references to predict cell types in unannotated datasets [11].

State-of-the-art automatic cell annotation methods perform well for most cell types but lack precision for rare ones. In some cases, simpler classifiers like Support Vector Machines (SVM) can outperform specialized methods [2]. Fully supervised methods are sensitive to batch effects, as they do not account for domain shifts between annotated and unannotated datasets. Batch effects may occur within reference datasets or between reference and query data. To address this, transductive semi-supervised learning can adapt models to the query dataset’s distribution [12]. For example, scNym combines a cell type classifier with a batch declassifier, using data augmentation and a domain-adaptation neural network (DANN) to correct for batch effects [13]. It performs data augmentation by creating hybrid cells from the annotated and unnanotated data following the MixMatch framework [14]. The authors also used a predictor combined with a domain-adaptation neural network [15] (DANN) to account for batch correction.

Integration and cell annotation are typically done separately, but hybrid models combine embedding learning with automatic annotation. These models aim to create a latent space that captures biological information for classifying cell types. For example, AutoClass combines an autoencoder and a classifier to learn a latent space from clustered annotated cells [16]. MARS uses an autoencoder to group cells around cell type landmarks and labels them based on their probability in a Gaussian-distributed latent space [17]. However, none of these models perform batch correction. scANVI addresses this by using a deep generative model with Bayesian inference to model counts in a latent space and incorporate cell type information for classification [4, 18]. Despite these advances, these models may struggle with low-abundance cell types. The highly imbalanced nature of single-cell data, with some rare cell types being crucial for pathogenesis and homeostasis, can hamper both integration and classification tasks.

This article introduces scMusketeers, a transductive cell annotation and integration model designed to address the imbalanced nature of single-cell data and highlight rare cell types. At its core, scMusketeers uses an auto-encoder to learn compact representations, reduce noise, and improve data recovery. Combining this with a classifier module enhances its classification ability, while an adversarial domain adaptation (DANN) module enables clustering analysis and batch effect removal. To improve performance on imbalanced datasets, scMusketeers incorporates focal loss, originally proposed by Lin et al in object detection [19]. We also tested a permutation technique, swapping cells of the same type during training, to enhance the embedding space. The combination of latent information reconstruction, DANN, and focal loss improved results. Our general modular framework allows to train and test an adaptive neural network architecture on various datasets in order to explore and optimize hyperparameters. We demonstrate scMusketeers’ robustness across 12 human organ datasets of varying sizes, cell types, and batch complexities. Its performance, using different module combinations, was compared with popular methods in single-cell integration and annotation. The three module of scMusketeers work together as *All for One* latent space, and this optimized latent space, which capture the essence of the single-cell data, resolves as *One for All* the initial challenges.

## 2 Results

### 2.1 Modular architecture of scMusketeers

scMusketeers is an automatic annotation model for single-cell data which can also be used as an integration tool. scMusketeers considers expression profiles coming from different batches, either annotated or not.

The core concept of scMusketeers is an all-in-one three-headed autoencoder which learns a low-dimensionality embedding in which cell type identity is reinforced. It is composed of three neural network modules (see Fig. 1A):

**Fig. 1.**
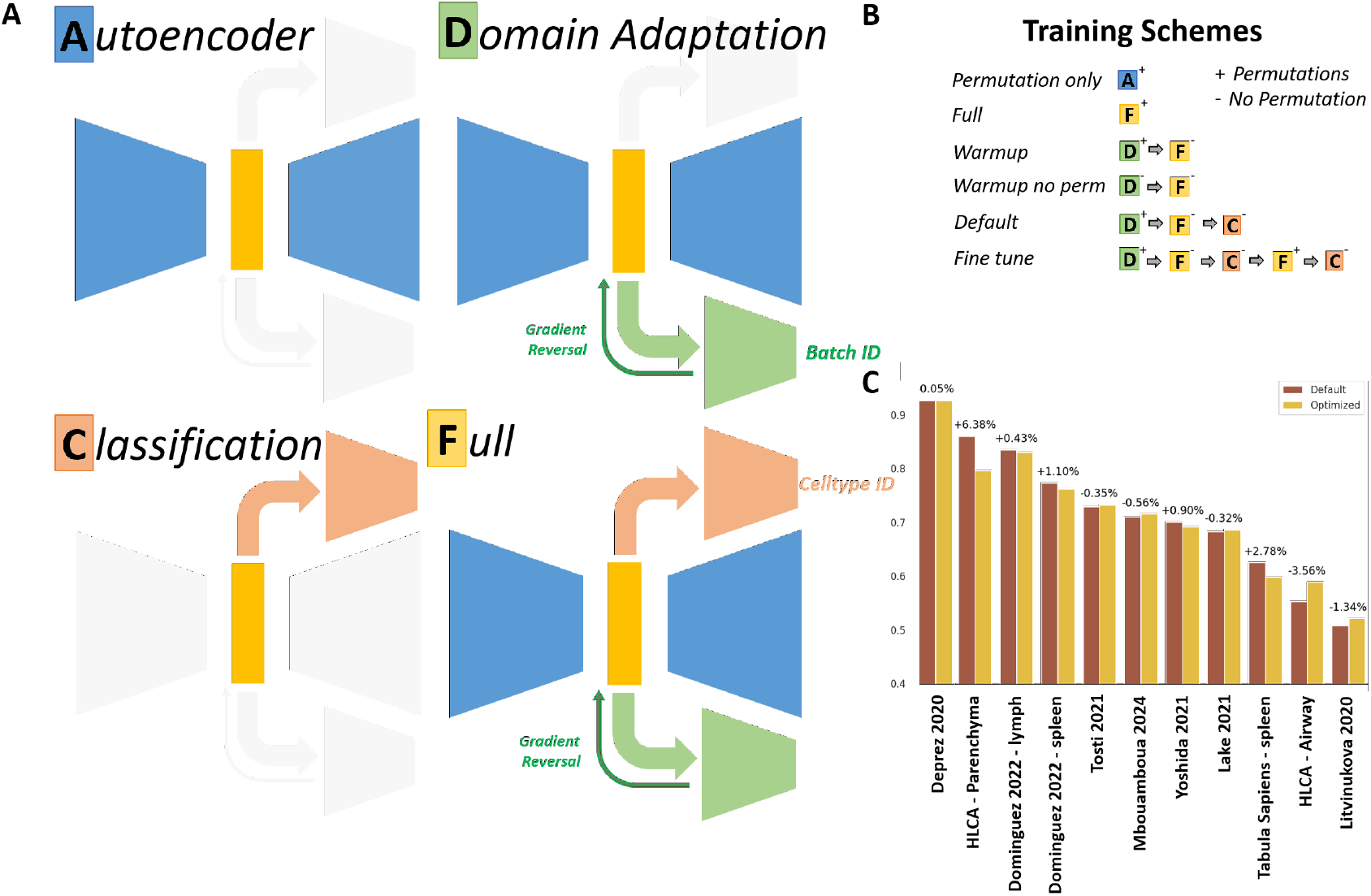
The scMusketeers model description. **A)** Architecture of the three headed model. Each head is optional. The autoencoder (blue) is a 2 layer autoencoder with MSE loss. The cell type classifier (red) is a standard MLP that aims at classifying cell types from the latent space, using a focal cross-entropy loss. The batch classifier (green) contributes negatively to the global loss via a gradient reversal layer that effectively acts as a declassifier. Four training modes are available, A (autoencoder only), D (Autoencoder + DANN), C (Classifier only) and F (All the modules at once) (see 9.5). **B)** Several training schemes were tested by alternating phases of training with each setup. Permutation can be either activated (+) or deactivated (-) at each phase for autoencoder reconstruction. The nomenclature showed here is used to describe the training scheme in this article. For example, the “Warmup” scheme will be written D+/F-. **C)** Barplots showing the mean balanced accuracies between the model with optimized hyperparameters and default hyperparameters (see Methods). Percentage number indicates the increase or decrease in accuracy between the default and optimized configuration.

- A two-layer autoencoder, which aims to build a latent representation of the expression space that uses mean-squared error (MSE) as a reconstruction loss.
- A cell type classifier plugged on the latent space layer in the form of a single-layer (MLP) that uses Focal Cross-Entropy (FCE) [19] as a prediction loss to improve classification of small cell types. This module makes the final cell type prediction.
- A batch declassifier plugged on the latent space layer in the form of a single-layer (MLP) that uses the gradient reversal technique proposed by Ganin & al [15] to perform domain adversarial training on neural network. We used Categorical Cross-Entropy (CCE) as a domain adaptation loss.

During backpropagation, the gradient of classification and domain adaptation loss flow into the encoder, therefore infusing the latent space with cell type and batch knowledge (see Supplementary Methods). The resulting latent space is then used as a low dimensional embedding in further analysis.

Because of its modular architecture, the training process can be divided in several steps where one or multiple modules are trained successively, as illustrated in Figure 1B. As part of those schemes, we also explored the idea of permutated reconstruction, where a cell is reconstructed as another cell from the same celltype by the autoencoder (see Fig. A2 A). The best training scheme was determined during hyperparameter tuning along with 11 other hyperparameters. From the hyperparameter tuning results found on 10 datasets, we were able to derive a default configuration for scMusketeers, which performed on par if not better than the optimized configuration on each dataset (see Fig. 1C). More details about training schemes, permutations and hyperparameter tuning are available in the Methods section.

We compared scMusketeers performances with 7 other models in 12 diverse datasets, using balanced accuracy as an evaluation metric (see Methods)

### 2.2 Transferring cell label across batches with scMusketeer

We evaluated scMusketeers’ ability to predict cell types on unknown batches, simulating the creation of a medium-large scale single-cell atlas from experiments processed over time. In this scenario, some of the data is annotated, and the goal is to transfer labels from earlier batches to newly produced ones, particularly in the presence of strong batch effects. Performance was assessed by training the model on a predefined training set and predicting on a test set. We repeated this process with five different train-validation splits, keeping or excluding entire batches (see Methods). Test and validation data were treated as unannotated by the models, with supervised methods not using them during training, and semi-supervised models using them as unlabeled data. Results were obtained using the default hyperparameters for scMusketeers, and cell prediction performance was measured using balanced accuracy.

scMusketeers outperformed other methods in cell type prediction on eight of twelve datasets: Tosti (+9%), Yoshida (+9%), Mbouamboua (+6%), Deprez (+4%), Dominguez-lymph node (+7%), Koenig (+3%), Litvinukova (+3%), and Lake (+8%) (see Fig. 2A). It performed similarly to UCE on the Dominguez-spleen dataset. scMusketeers was better when integrating datasets rather than batches in the two HLCA datasets, where it was matched only by Celltypist and Harmony in the HLCA-airway and by scANVI in the HLCA-parenchyma. Notably, scMusketeers showed consistent performance, unlike scANVI or Harmony, which had more variable results. For example, scANVI was best in the HLCA-parenchyma study, and Harmony in the HLCA-airway study, but both underperformed on the Deprez and Lake datasets. On the Tabula Sapiens dataset, all models performed similarly, with no clear standout.

**Fig. 2.**
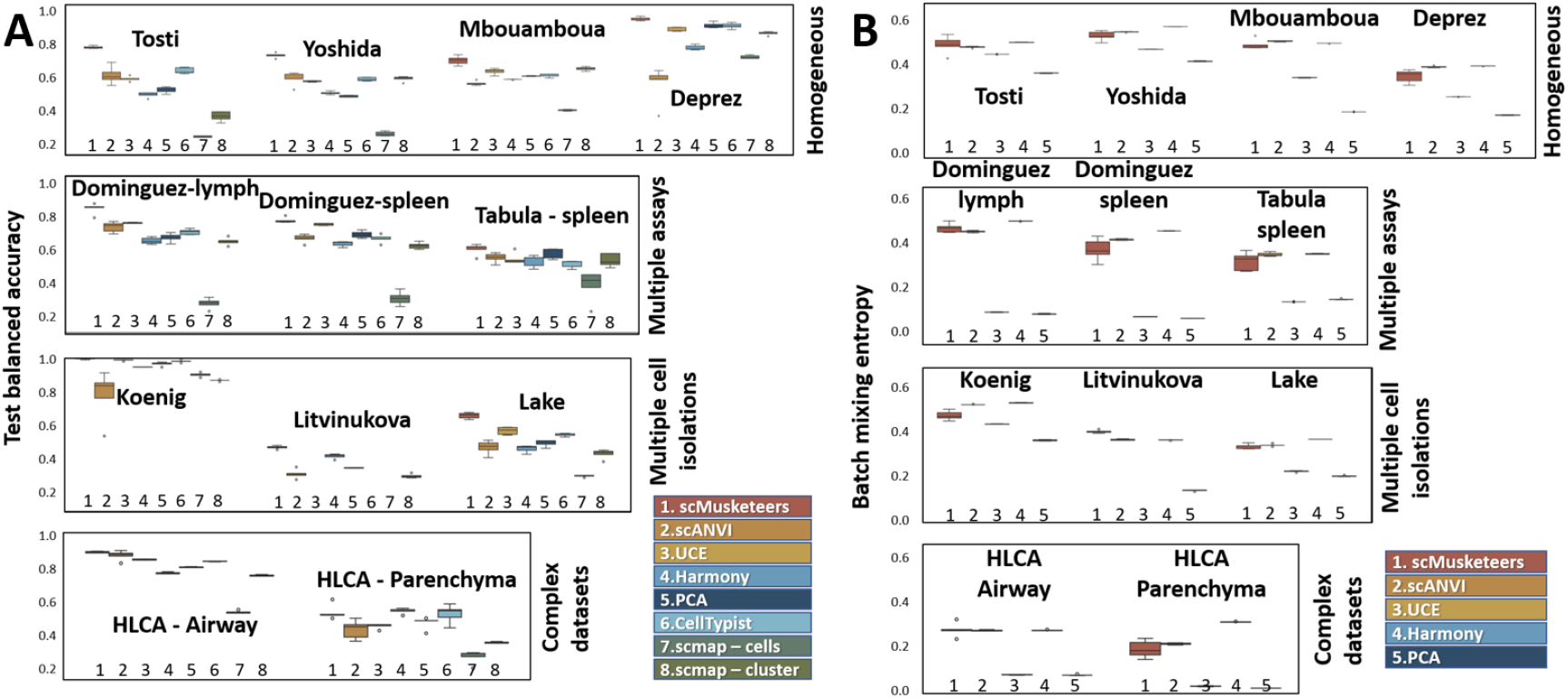
**A)** Balanced accuracy on the test split for each dataset. Each training was performed on 5 different train/validation splits. Datasets on the same row belong to the same source of batch effect: “Homogeneous” if samples came from similar sampling and analysis conditions, “Multiple Assays” if different assays were used in the dataset, “Multiple Cell Isolations” if the dataset mixed single cell and single-nuclei isolations, “Complex Datasets” if the batches were different datasets themselves. **B)** Batch mixing entropy computed for each dataset on the same experiments as Fig. 1. Batch mixing entropy was computed on the entire dataset including the train and val splits, to observe the good mixing of the query dataset with the reference.

### 2.3 Batch removal capability of scMusketeer

We assessed the batch removal capability of each model using the batch mixing entropy (BME) score. scMusketeers showed BME values similar to integration tools like scANVI and Harmony, indicating comparable batch removal abilities (see Fig. 2B). A high BME generally suggests good batch removal in low-dimensional representations, even though it does not guarantee preservation of high-dimensional structures. scMusketeers performed on par with common integration methods. UCE, despite aiming for universal representation, showed lower scores, performing between integration methods and PCA when batch effects were mild. However, with more severe batch effects (e.g., multiple assays or complex datasets), UCE’s BME scores matched those of PCA, which does not correct batch effects.

The variance in batch entropy was high in scMusketeers, particularly when the batch effect was associated with different sequencing assays. However, this variability was not observed in prediction accuracies, where scMusketeers’ variance was similar to that of other models. Thus, there was no clear relationship between batch removal effectiveness and predictive power in scMusketeer.

### 2.4 scMusketeers is highly effective for detecting rare cell-states

We assessed the ability of various models to detect rare cell types, a challenging task due to the imbalance in single-cell datasets, which often biases models towards more abundant classes. Detecting rare cell types is crucial, as they can play significant physiological roles, such as pulmonary ionocytes, which, despite being a small fraction of airway cells, are linked to cystic fibrosis (CF) manifestations [20–22]. To evaluate rare cell type detection, we defined two levels of low abundance: (1) less than 0.1% of the dataset (tiny) and (2) between 0.1% and 1% (rare). We set a detection threshold with an accuracy score above 20%, reasoning that identifying rare cell types, even with low accuracy, is sufficient for single-cell atlas construction, with further refinement possible through manual annotation.

Figure 3A and 3B show the average number of rare (<1%) and tiny (<0.1%) cell types identified per dataset by each model (5 repetitions). Most models struggled with tiny cell types, but scMusketeers outperformed all methods, except scmap-cluster, across datasets. For example, scMusketeers identified more cell types in the Lake and Yoshida datasets and was one of the only methods (along with scmap-cluster) to detect tiny cell types in the Tosti dataset. Annotation-specific methods like Celltypist and scmap-cluster generally outperformed embedding methods like scANVI and Harmony, except for UCE, which ranked third overall.

**Fig. 3.**
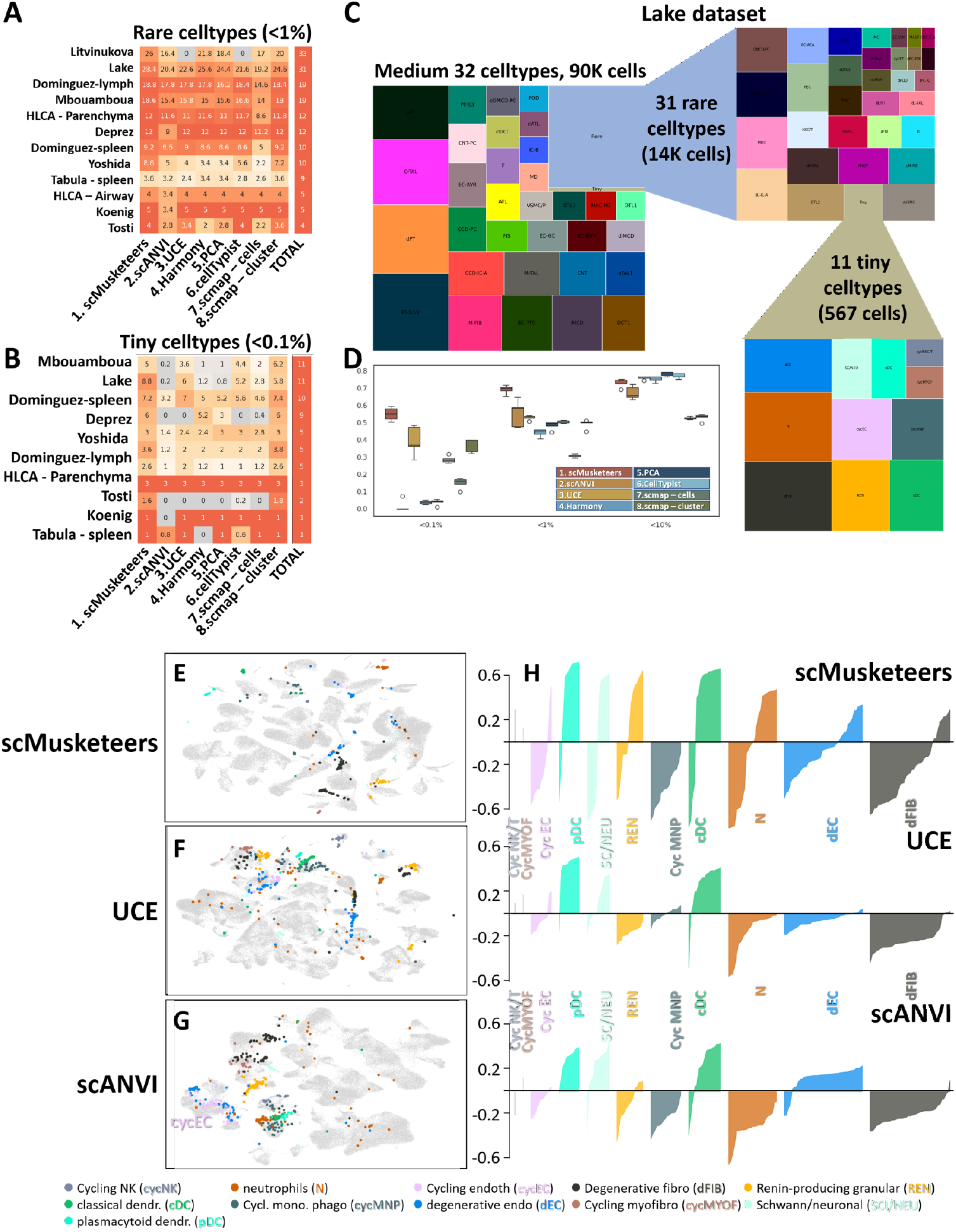
**A)** Number of tiny (left panel) and rare (right panel) cell types identified with an accuracy superior to 20% by the best prediction for each model. The right column of each table (Total) represents the total number of such cell types in each dataset. **B)** Treemap representing the class imbalance in the Lake dataset, broken down into the medium, rare and tiny cell types. The size of the squares represent the total proportion of this cell type in the dataset. **C)** Test balanced accuracy according to the cell type size in the Lake 2021 dataset. Cell types are divided in four bins depending on how much of the dataset they represent (<0.1%, <1%, <10% and >10%) and balanced accuracy are computed independently on each bin. Each column corresponds to a different method (see legend). UMAP representations derived from **A)** scMusketeers, **B)** UCE and **C)** scANVI of the Lake dataset. Each UMAP was obtained from the latent space computed by the corresponding method. Only cells belonging to the tiny cell types (<0.1%) were highlighted. **D)** Silhouette score profiles on the tiny cell types of the lake dataset computed in the latent space of each method. A score close to 1 corresponded to cells which located within their own cluster, a score close to -1 correspond to cells overlapping with other clusters rather than their own.

For rare cell types (i.e. <1%), detection improved across all models. scMusketeers outperformed all methods, including scmap-cluster, particularly on datasets with many rare cell types, such as Lake (+2.8) and Litvinukova (+6).

We further explored these findings in the Lake dataset, which describes an atlas of the kidney under different physiopathological conditions. In this dataset, we considered the subset that corresponds to the healthy donors. It includes 74 cell types and/or sub-types. Those sub-types correspond to different cell states defined by the author as normal, cycling, adaptive or degenerative. Figure 3C shows the severe class imbalance, with 11 cell types representing less than 0.5% of the dataset and 32 cell types representing more than 90% of the dataset. Contrary to other datasets, no individual cell type was overwhelmingly represented. The annotation task with this dataset was complicated by the high number of closely related cell types/states.

scMusketeers was slightly better than the other models in terms of global balanced accuracy (Fig. 2A). Figure 3D reveals that this better performance was particularly due to the tiny cell types for which other models exhibited lower accuracies. Similar observations were made with the other datasets (see Fig. A3-A4).

Notably, we also observed that the high detection rate of tiny cell types by scmapcluster seen in Figure. 3A-B was balanced by a poorer performance for the annotation of larger cell types (see Fig. A14). On the Lake dataset, scmap-cluster was less accurate on larger cell types. By contrast, scMusketeers maintained a high level of detection across all cell types, regardless of their respective sizes. The same can be said for UCE, although with an overall lower accuracy compared to scMusketeer.

Considering possible contributions of rare cell types to pathologies, and their more difficult identification by machine learning methods that are more biased towards broader classes, scMusketeers provides a useful framework for such identifications.

### 2.5 scMusketeers embedding space accurately captures cell type imbalance

The better detection of rare cell types is arguably a consequence of using a focal loss, which improves detection of smaller classes. Besides, this property permeates to the latent space, such that scMusketeers builds a latent space in which rare cell types are better segregated from larger populations. The Lake dataset contains eleven particularly challenging tiny cell types :

- Two rare cell types of the kidney: renin-producing granular cells (REN) and Schwann/neuronal (SCI/NEU) cells.
- Two degenerative cell states, i.e. degenerative endothelial cells (dEC) and degenerative fibroblasts (dFIB), which are mostly present in pathological cases and less abundant in healthy donors.
- Four cycling cell states for endothelial Cells (cycEC), myofibroblasts (cycMYOF), mononuclear phagocytes (cycMNP) and NK cells (cycNK).
- Three myeloid cell types: neutrophils (N), classical dendritic Cells (cDC) and plasmacytoid dendritic Cells (pDC)

Myeloid cell types can be difficult to characterize in single cell studies. Moreover, the cycling and degenerative populations are also hard to detect, given that they correspond to cellular states that alter existing cell types.

In Figure 3E-H, we compared the three Deep Learning methods scMusketeers, UCE and scANVI which provide a batch-free embedding. The UMAP representation derived from scMusketeers grouped those rare cell types/state in separate clusters, isolating them from their respective “mother” cell types/states (Fig. 3E). This separation was less clear for UCE and scANVI, in which those cell types were mixed together with neighboring cell types, thus explaining why they were not detected by these models. Instead, identification of distinct clusters in the scMusketeers latent space allowed their detection.

The observation made on the UMAP was confirmed on the whole data set by computing silhouette scores for every tiny cell in the test set. As we evaluated the latent space rather than the predictions, we used the true labels of the cells rather than their predicted label to compute the silhouettes (see Methods). A positive score indicates cells grouping closely to each others in the same cell type. A high positive value indicates a better compacity of the corresponding cluster. The silhouette profiles in Figure 3H indicate that pDC, cDC, sc/NEU, cycMYOF, cycNK/T were separated equally well by all methods. scMusketeers managed to partially isolate dFIB, dEC, N and REN in the test set. Depending of the cell type, 25-50% of the cells showed high positive silhouette score, indicating their grouping together with other cells from the same cell type in the reference. This was clearly not the case for the other models apart from the dEC cells accurately identified by scANVI (Fig. 3H and Fig. A13).

These silhouette profiles are consistent with the observed prediction accuracies of each cell type made by scMusketeers and UCE, meaning that a better discrimination of the cell types leads to a better detection by the models (see confusion matrix in Fig. A13). For all three models, misannotations were not totally inconsistent. Instead, they rather identified a similar cell type or the initial cell type from which the cell state was derived from. For example, scANVI confused cycling NK and fibroblasts as regular NK and fibroblasts, respectively, and all three models confused the cycMNP as M2 macrophages. Surprisingly, while scANVI managed to isolate 4 of those cell types, Fig. 3D and Fig. A13 shows that the model did not detect them. This might come from the scANVI model making predictions using its internal encoding of the cell type identity as a latent vector, rather than using a straightforward classifier for annotation.

### 2.6 Annotation transfer from partially labelled datasets

We then assessed scMusketeers in the task of seed labelling, that is, predicting cell types when trained on only a small portion of a dataset across batches. To do so, we considered two setups: (i) In the first setup, we selected a percentage of the training set to use as annotated training data and the rest of the training set to use as validation data. We ran this experiment using 5%, 10%, 50% and 90% of the training set. (ii) In the second setup, we selected a fixed number of cells from each cell type in the training set to use as annotated training data. The rest of the cells was used as validation data. We ran this experiment with 5, 10, 50, 100, 500 cells of each cell type in the training set. When a cell type was smaller than this number, we selected the whole cell type.

The first setup therefore maintains the cell type proportions while the second one is supposed to mitigate class imbalance, by having the same number of cells from each cell type to train on.

Results show that scMusketeers generally outperformed the other methods when trained on the same quantity of data (Fig. A9 to A12). scMusketeers performance was assessed by considering the minimal amount of training data that was necessary for scMusketeers to outperform the other models that were trained with the maximal amount of data. Fig. 4A corresponds to setup (i), and represents the percentage of the data at which point scMusketeers outperformed other models that were trained on 90% of the dataset. scMusketeers usually needed 50% or less of the data to achieve the same performance as other models trained with 90% of the data. scmap, PCA and Harmony showed lower performances at this task. On most datasets, scMusketeers only needed 5% or 10% of the data to perform better than these three models, even though they were trained on the whole dataset (90%). scMusketeers also outperformed Celltypist, scANVI, UCE, but in these cases, it often needed 50% of the training data to achieve similar results on these methods when they were trained on the whole dataset. scMusketeers seems especially good on the Lake dataset, where it outperforms every models but UCE with only 10% or less of the dataset. We also saw that other methods were not performing well on the same datasets : UCE had poor performances on Tosti and HLCA-Airway compared to scANVI and Celltypist. On the contrary, UCE did better on Mbouamboua where scANVI and Celltypist were outperformed with 10% or less training data.

**Fig. 4.**
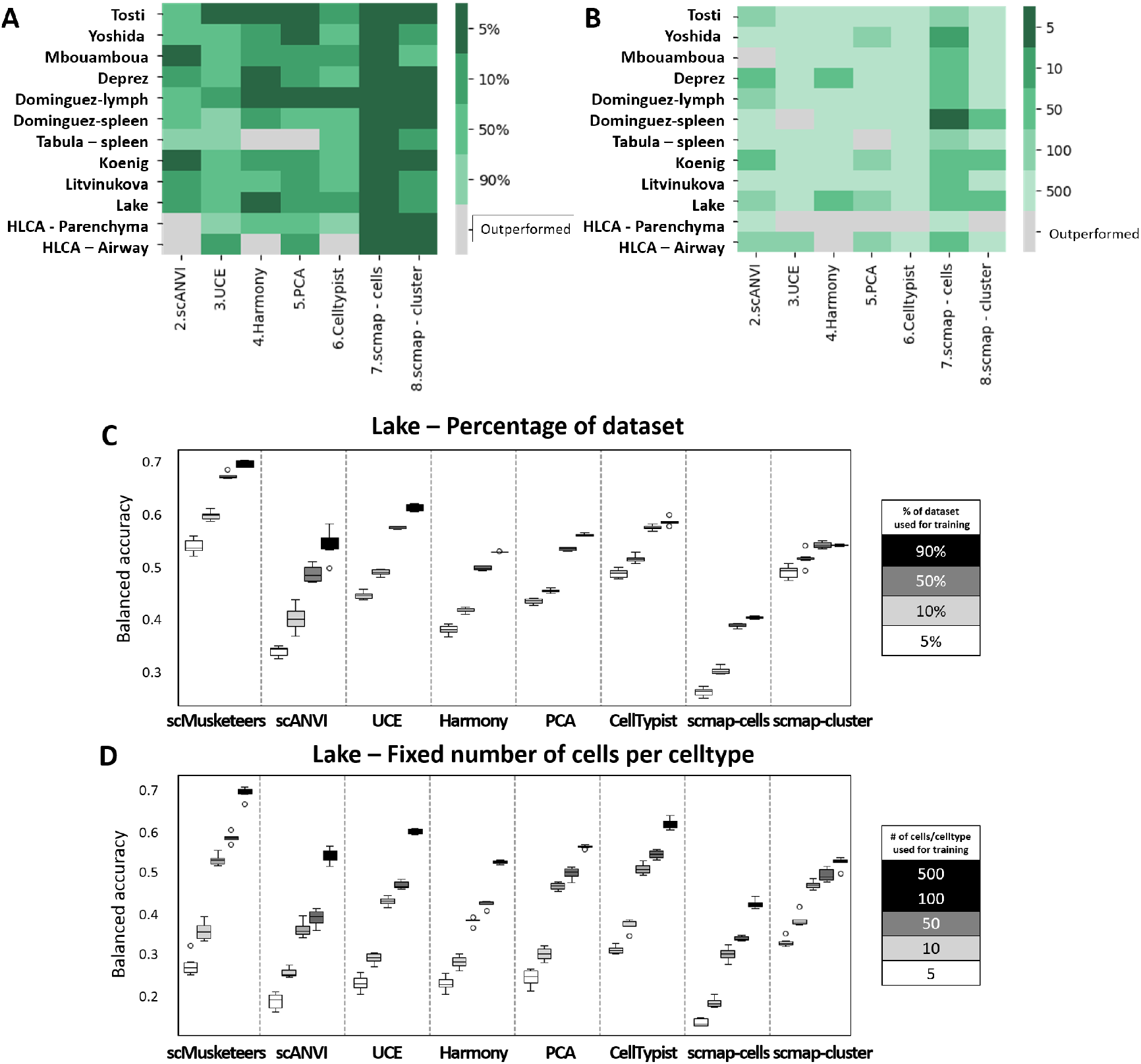
**A)** Heatmap representing the percentage of the dataset used for training at which scMusketeers outperforms the other models trained on 90% of the dataset **B)** Heatmap representing the number of cells per cell type selected for training at which scMusketeers outperforms the other models trained on 500 cells per cell types. **C)** Balanced accuracy on the Lake dataset depending on the percentages of the dataset or **D)** the number of cells per celltype used for training.

Figure 4B corresponds to setup (ii), and represents the number of cells per cell types (cells/ct) at which point scMusketeers outperformed other models trained on 500 cells/ct. Notably, training on a fixed number of cells per cell type (cells/ct) yields very different results. For the most part, scMusketeers is better than other models when trained on the same number of cells but it is rarely better with less cells (Fig. 4B and Figure A9-A10). scMusketeers consistently outperformed scmap-cells and scANVI with less annotated cells. Surprisingly scANVI was the only model to outperform scMusketeers on Mbouamboua when trained on 500 cells/ct but it was completely outclassed by scMusketeers in the previous experiment. There, scMusketeers needed only 5% of the dataset to beat scANVI. scMusketeers top performances still appear to be on the Lake dataset, where it outperformed every models but Celltypist and UCE with less cells, needing only 100 and sometimes 50 cells/ct to outperform.

On Figure 4C-4D representing balanced accuracies on the Lake dataset, we see that keeping the cell type proportions seemingly leads to better results in terms of balanced accuracy. The dataset is composed of approximately 100 000 cells for 74 cell types which means that 5% of the dataset can be obtained by taking 100 cells by cell type, considering that some cell types have less than 100 cells. However, scMusketeers achieved better results when training on 5% of the dataset while keeping cell type proportions intact rather than taking 100 cells of every cell types. This is surprising because we expected that equalizing the classes would result in a better prediction of the less represented ones and therefore a better balanced accuracy.

### 2.7 cell type prediction on spatial transcriptomic

We finally evaluated how scMusketeers was transferring labels from a reference dataset to a spatial transcriptomics one. This type of experiments has several special features: the batch effect is stronger than in single-cell RNAseq experiments, capture efficiency is low and only a fraction of the genes is analyzed. It was therefore interesting to assess the performances of scMusketeers under these more extreme conditions. We tested a public dataset generated with the Xenium technology from 10X Genomics, accessible on the company website [23], corresponding to an healthy lung parenchyma. The dataset captures the expression of 390 genes across roughly 200,000 cells. cell types were transferred after a training of every model on the parenchyma compartment of the HLCA [24], then predicting on the spatial dataset. Training was performed on a coarse level of annotation, given that the experiment only analyzed 390 distinct genes.

In the absence of curated cell types for this dataset, a qualitative appreciation of the different performances was done.

All models except scANVI and scMusketeers provided aberrant cell type proportions, 39% of T cells for Harmony, 53% of Fibroblasts for the PCA and 50% of Macrophages for Celltypist as shown in Figure 5A. Most likely, those models did not adapt well to the strong shift in distribution. Only scMusketeers and scANVI identified cell type proportions coherent with the expected composition of the lung parenchyma, which led us to further explore those 2 tools. Notably, scMusketeers identified 12% more Fibroblasts compared to scANVI. On the opposite, some cell types were identified in very small quantities by scMusketeers in particular venous and lymphatic endothelial cells, as well as bronchial epithelial cells all composing less than 1% of the total tissue.

**Fig. 5.**
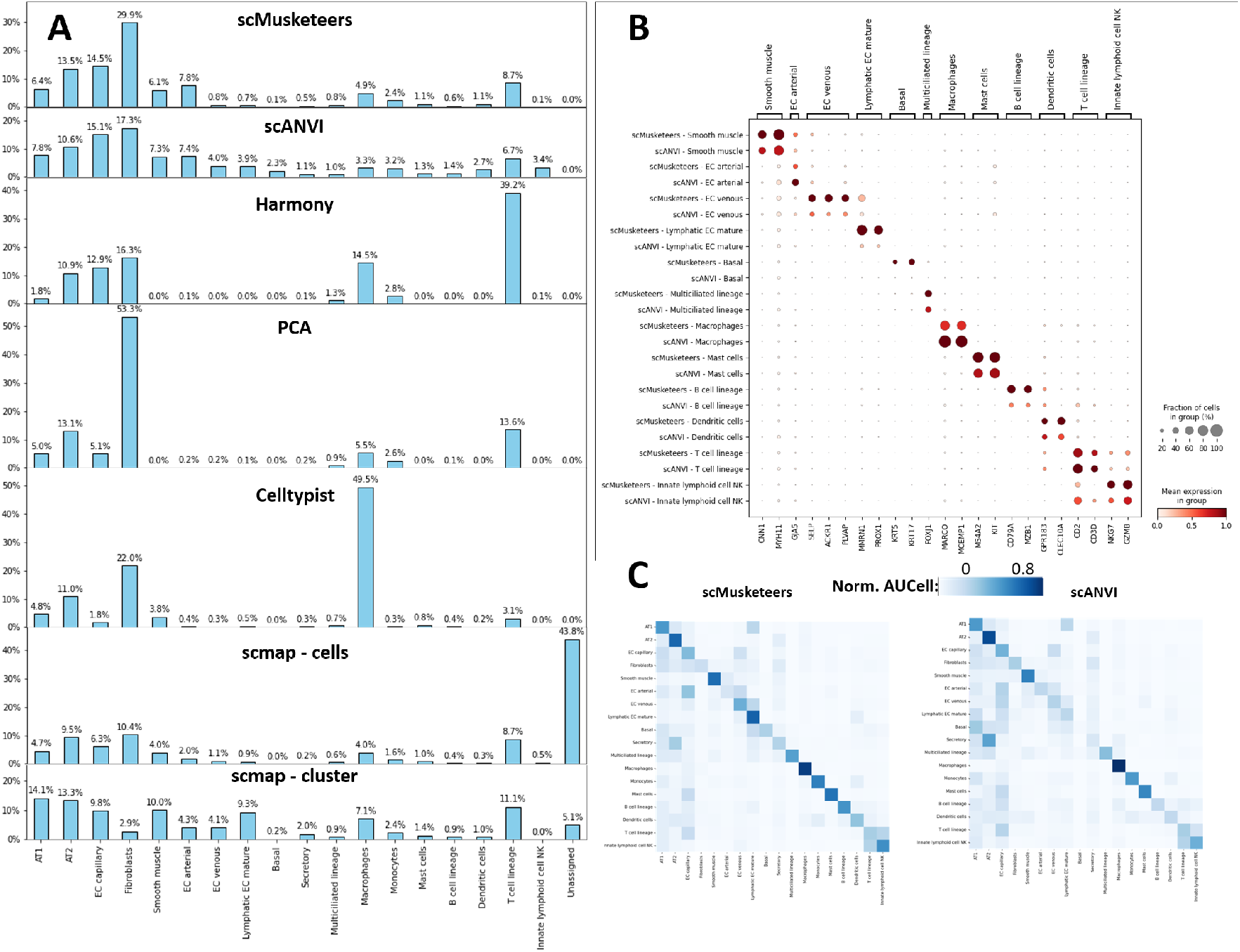
**A)** Proportion of cell types predicted by the different models in the Xenium lung dataset. The “Unassigned” category corresponds to the rejection option only available for scmap. **B)** Dotplot of relevant marker genes for the scMusketeers and scANVI annotations. the size of each point indicates the percentage of cells from this cell type expressing the gene. Expression values are scaled between 0 and 1. **C)** Heatmaps of normalized AUCell scores computed from reference marker genes. Y axis corresponds to the predicted cell types and X axis to the gene signatures.

We further investigated those discrepancies by looking at known marker genes in cell types for which results differed between the two annotations (see Fig. 5B). Higher fractions of endothelial venous and lymphatic cells, as well as Basal cells, B cells, Dendritic cells, and NK cells were found by scMusketeers, compared to the scANVI annotation. In particular, cell types that represent around 1% of the dataset were identified in lesser amounts by scMusketeer. This came along with a better score in term of marker genes, thus suggesting a higher precision of scMusketeers for those small cell types. Conversely, scANVI exhibited slightly better results for macrophages and endothelial arterial cells. Known markers were used to compute cell type signatures using the AUCell tool [25]. Results are summarized in Figure 5C. Those cell type scores further reinforce those observations, as the aformentioned cell types exhibit weaker scores for scANVI compared to scMusketeers, especially in the case of the Basal and immune cells.

To further evaluate the quality of those annotations, we detailed the cellular profiles found in three distinct parts of the lung parenchyma, i.e. vascular, bronchial and alveolar niches (see Fig. 6A-C). Results were similar in both models and they appeared coherent with their expected compositions of those structures. In the bronchial niche, both models properly identified basal, secretory and multiciliated cells, as expected. Those were surrounded by a few fibroblasts and smooth muscle cells. A very consistent spatial organisation of those cells was observed with both models (Fig. 6D-F). Notably, scMusketeers only identified basal cells in the specific bronchial regions of the tissue, whereas scANVI also found basal cells across the parenchyma in alveolar regions.

**Fig. 6.**
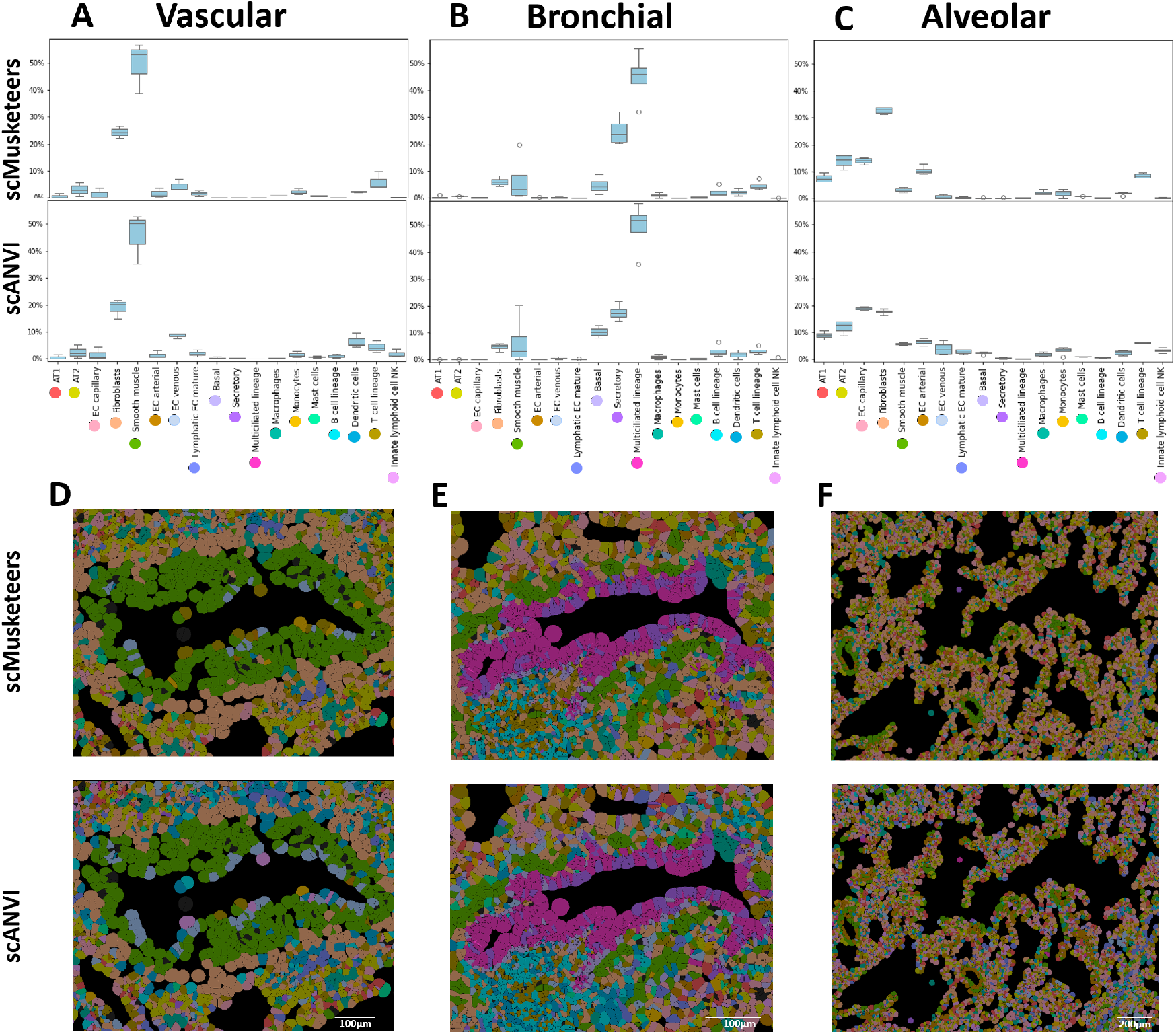
Proportion of cell types in **A)** vascular, **B)** alveolar and **C)** bronchial niches of the parenchyma. Each boxplots were made considering 4 different regions of each type. Spatial representation of cell type organisation identified by scMusketeers and scANVI in an bronchial niche **D)**, a vascular niche **E)** and an alveolar niche **F)**

The vascular niche displays 2 concentric layers of fibroblasts and smooth muscle, with an inner layer of endothelial cells and a few immune cells. Figures 6D shows that the organisation of the cell types is consistent with both annotations, with stromal cells surrounding the blood vessel, endothelial cells lining the internal surface of the vessel and T cells and macrophages identified in the lumen of the vessels. scANVI identified more clearly the endothelium in the vasculature compared to scMusketeers, but this is probably explained in part by the fact that scANVI identified a larger number of endothelial cells (see Fig. 5A). This favored a better recall rather than precision for this cell type. This leads to better delineation of the endothelium in the vasculature but a worst marker gene signal in endothelial cells which corresponds to a higher number of false positives.

The alveolar niche showed very similar profiles of AT1, AT2 and capillary cells across both models, as illustrated in Figure 6F that shows the organisation of an alveoli.

Collectively, these results posit scMusketeers as an appropriate tool to annotate a spatial transcriptomics dataset. Given the satisfactory results already provided by scANVI, we recommend to use the intersections between the two tools as a core annotation of the cells before any further annotation. It could also be a good starting point to design tools specifically designed for this task in the future.

## 3 Discussion

We developped scMusketeers, an automatic annotation tool which builds a cell type-informed embedding space. To do so, we used a modular architecture composed of an autoencoder, a cell type classifier and a batch discriminator. We noticed that the use of focal loss for the cell type classifier improved performances to detect rare cell types without compromising the annotation of the remaining dataset. We also tested for the first time the use of permutated reconstruction, which yielded marginal improvements but we believe should be explored further.

We benchmarked scMusketeers against 7 popular automatic annotation methods on 12 datasets, representing a large variety of human, organs, sizes, number of cell types and batch effects. An hyperparameter tuning on 10 of the smaller datasets allowed to derive a default configuration which generalized well to the larger ones.

scMusketeers significantly outperformed the other methods on most datasets, indicating its generalization power. We also qualitatively assessed scMusketeers in the task of label transfer from scRNAseq to spatial transcriptomics, where it yielded convincing annotation with regards to spatial structures and marker genes expression.

Notably, every classification results in this article were evaluated using balanced accuracy as well as binned balanced accuracy depending on cell type size. While this approach is uncommon in the scRNAseq litterature, we believe it is necessary to take into account the severe class imbalances existing in single-cell datasets. scMusketeers demonstrated superior performances, particularly in accurately detecting rare cell types and separating them from similar cells in its embedding space. This model also exhibited a notable advantage when trained on a small quantity of data compared to other models. In the many single-cell RNAseq experiments in which datasets are annotated “en masse”, this property is not necessary useful. This could however open the door to new ways of annotation based on curricul’hum or active learning, by iteratively feeding a few proportion of expertly annotated cells to the model [26–28].

There are nowadays different strategies to annotate a single cell dataset, and it is likely that each will come with its own pro and cons. scMusketeers good performances are counterbalanced by a few limitations that should be addressed in the future.

First, we built scMusketeers by considering the current state of the art in single-cell datasets which generally integrate several batches. As a consequence, scMusketeers performs poorly if there is a single annotated batch in the training data. scMusketeers does not explicitly model batch effect and batch correction is performed through the combined effort of the DANN and classifier as well as the permutations. scMusketeers needs several annotated batches to learn a generalized batch free representation of the data.

Second, scMusketeers does not currently allow cell type discovery. Some automatic annotation model have an option to identify cell types not present in the training dataset. To do so, MARS defines a set of landmarks in an unsupervised way in the query data. One shortcoming of this method is that MARS requires to know the exact number of cell types in the query for an accurate prediction. OnClass leverages external data from the CellOntology database to build an embedding space, and learns a function to map RNAseq data in this space [29, 30]. OnClass is theoretically able to identify every cell type that exists in CellOntology. A more modest approach consist in identifying every unseen data as one “unknown” or “novel” type. scNym and scBert perform this identification using a confidence threshold on the output probabilities of the classifier [13, 31]. Especially, scNym combines this with temperature scaling to smooth out the output probabilities. This could be implemented in the current scMusketeers model quite straightforwardly.

Third, we only ran a limited number of 50 trials per dataset on a set of 11 optimized hyperparameters. While this yielded encouragingly good results, one could argue that this was insufficient to explore such a large hyperparameter space. Ideally, further exploration should be performed. We overall advocate for a better hyperparameter search in the domain of single-cell automatic annotation as most published model are often made available with arbitrary default parameters.

A significant observation in our study was the substantial variation in average performance across all datasets, irrespective of the model employed. This suggests that inherent dataset characteristics, such as complexity and data quality, significantly influence model performance and the cell annotation process. Despite the development of many quality control metrics, and best practices to ensure the quality of embedding spaces and automatic annotations [1, 7, 32, 33], cell annotation still relies on extensive human curation. Given our observations, we believe that the cell annotation process should be used as a proxy to better define the quality of single-cell datasets. By analyzing the performance gap across datasets and identifying the cell annotation metrics correlated with these performance variations, we could gain deeper insights into the factors that contribute to the quality of a single-cell RNA-seq dataset.

## 4 Availability and requirements

scMusketeers is available for use on PyPi. Documentation can be found at https://sc-musketeers.readthedocs.io/en/latest/. scMusketeers code can be downloaded in https://github.com/AntoineCollin/scMusketeers. A tutorial can be found here.

## 5 Acknowledgements

The authors are grateful to the Université Côte d’Azur’s Center for High-Performance Computing (OPAL infrastructure) for providing resources and support.

## 7 Authors’ contributions

Conception and design: A.C., F.P; Benchmark implementation: A.C, M.F, S.P.; Analysis: A.C.; interpretation and validation: A.C., F.P., C.B., P.B.; Drafting of the manuscript: A.C., P.B.; Package publishing: C.B, A.C; Manuscript review and edition: all authors; Funding and supervision: F.P, P.B.

## 9 Methods

### 9.1 Testing cell permutation in scMusketeers training process

To enhance cell type identity and reduce batch effects, we explored permutated reconstruction in the autoencoder. Unlike traditional autoencoders, which reconstruct the original input from the latent space, this permutated reconstruction model reconstructs the expression profile of a randomly selected cell from the same cell type rather than the original cell (Fig. A2 A). This is related to the Centroid-Autoencoder reconstruction proposed by Ghosh et al, which used instead the centroid of the input’s class [40]. Our approach performs intra-cell type permutations during training, matching each cell with a different one from the same type, strengthening cell type grouping in the latent space. Permutation has two benefits: (1) it increases training pairings, boosting small populations, and (2) it helps mitigate batch effects by performing permutations across batches. Unannotated cells were assigned initial pseudolabels using an SVM trained on annotated cells in the PCA space, with the model’s predictions serving as pseudolabels after the first training round.

### 9.2 HYPERPARAMETER tuning and training scheme selection

Hyperparameters were optimized using a bayesian search on each dataset in 50 trials on 10 parameters .For each dataset, we selected the run with the highest balanced accuracy. Since hyperparameter search is very time and ressource consuming, we computed a default configuration derived from the best configurations of each dataset (see Methods). The two largest datasets, i.e. Litvinukova and HLCA-Parenchyma, were not included to compute the default configuration, and were instead used to independently assess the quality of this configuration. This avoided an expensive hyperparameter optimization on 2 very large datasets. We kept the top 5 performing configurations for the remaining 10 datasets. The default configuration was obtained from those 55 configurations by taking the average or log average of numerical parameters and the most frequent parameters in the case of boolean or categorical parameters. The only exception is the number of HVG that we set at 3000, in the sake of fairness with other models. Notably, the default configuration performed on par if not better compared to (see1 C) the optimized versions, indicating the robustness of the default parameters found by our approach. This is consistent with the observations made on the Litvinukova and HLCA-parenchyma datasets, which gave good results, although they were not taken into account to calculate the default configuration. This suggests that the hyperparameter optimization can be performed on smaller datasets and then be extended to larger ones, in a less-intensive computational set-up.

Different training schemes were explored to optimize adversarial training of the whole model. We separated the training process in several steps, training one or multiple modules of the model successively, with or without permutations (Fig. 1B). Four training schemes were tested during hyperparameters tuning, illustrated in Fig 1B : (1) D+/F+, (2) D+/F-, (3) D-/F-, (4) D+/F-/C, in which D uses the autoencoder plus the DANN, C uses only the classifier, F uses the autoencoder, the classifier and the DANN. +/-indicates the use of permutations or not. All of these schemes include a warmup step, which consists in a simultaneous training of the autoencoder with or without permutations together with the batch declassifier. This initializes the full training with weights that already corrects for batch effect while capturing cell type identity via permutations. The best scheme during hyperparameter search ended up being D+/F-/C, that is a warmup, followed by a training of the full model, then a refining of the classifier. Most results on scMusketeers, including the default setup, were generated using this scheme.

We only tested four training schemes in order to limit the complexity of the hyper-parameter search. Additionally, we tested other candidate schemes with the default hyperparameter configuration to have an assessment of this default training scheme on five datasets (see Fig. A1). For every training scheme with permutation, we evaluated its counterpart without permutation. Figure A1 shows that better results could have been obtained with another scheme. Notably, the D+/F-/C/F+/C exhibits top BME scores and balanced accuracy scores with low variance. In this scheme, new pseudolabels are computed for the unannotated cells after the first classifier training. Those labels are used for the permutations of the following F+ step. Although marginal gains were observed when using the permutation on three datasets, further work is still necessary to explore the full potential of such a training setup (see Fig. A2 B). For the sake of comparison, we implemented two scMusketeers training schemes replicating scNym and Autoclass architectures. We did not add directly these tools in our benchmark because their corresponding GitHub repository are no longer maintained. Our workflow allowed to replicate similar architectures, improved by the use of the FCE prediction loss and our optimized hyperparameters. Both architectures performed poorly on 2 datasets (i.e. Tabula Sapiens and Mbouamboua) but were on par with the other architectures on the three other ones.

### 9.3 Benchmarking scMusketeers with other automatic annotation methods

We benchmarked scMusketeers against five state-of-the-art integration and annotation tools, based on deep or machine learning approaches. we used two integration tools, UCE [6] and Harmony [5], two annotation tools, scmap [41] and Celltypist [11] as well as one hybrid tool, scANVI [4]. When only an embedding was available in the case of UCE and Harmony, cell types were predicted after training a SVM in the embedded space. We also included a baseline tool in the benchmark consisting of a PCA followed by a SVM trained on the PCA space.

### 9.4 Balanced accuracy instead of accuracy

We assessed the quality of our predictions by measuring balanced accuracy, by weighting the accuracy of each cell by the inverse of its cell type size.This is important because single-cell datasets are highly imbalanced in cell type size and performance should always take into account this asymmetry [42]. Prediction of rare cell types is a key question in cell type prediction. To evaluate this task, we also computed the balanced accuracy on 4 cell type subsets according to their prevalence in the dataset (tiny [<0.1%], rare [0.1% - 1%], medium [1% - 10%], and large [>10%]).

scMusketeers not only transfers cell types but as an autoencoder it also produces an informative embedding space. Integration performance during benchmarking of single-cell integration tools was assessed using the batch entropy mixing metrics described by Luecken & al [1] (see Methods).

### 9.5 Datasets selection for the benchmark

We used datasets of various size and complexity coming from distinct human organs to assess the adaptation and scalability of our method (see Supplementary Table 1). They account for different levels of single-cell dataset complexity. Every one of them includes biological and technical replicates inducing a batch effect in the integrated dataset. Selected datasets differed by three criteria : number of cells (ranging from 10K to 450K), number of cell types (from 14 to 74) and number of batches (from 3 to 35). In addition, these datasets represent four types of batch effects with varying degrees of complexity :

- Homogeneous, when batches were generated with identical technology and preparation of the cell suspension. This is the case of: Mbouamboua (in preparation), Deprez [39], Tosti [34], and Yoshida [35].
- Multiple Assays, when batches came from different single-cell RNA sequencing technologies, such as different versions of 10X protocols, or use of Smart-seq2. This is the case of Dominguez [11], which we separated in a spleen and a lymph node datasets. Both also use different versions of the 10X single cell 3’-RNA sequencing protocol. Splenic cells from Tabula Sapiens [38] combines samples from 10X and Smart-seq2.
- Multiple cell isolations, when batches contained different types of single-cell RNA sequencing approaches, i.e.such as single cell versus single nuclei analyses (nuc-seq). This is the case of Koenig [36], Lake [43] and Litvinukova [37]. The latter also mixes different versions of 10X assays.
- Complex datasets, which are composed of multiple datasets, combined to create an integrated atlas. We consider datasets as batches per se. We assessed label transfer in the HLCA [24] in which we separated the Parenchyma and the Airway compartments.

The test sets were selected randomly while making sure that they preserve the same cell type proportions than the initial dataset. We performed 5-fold validation on three different train-test splits with every model (except scMusketeer) for each dataset. The test sets yielding the overall higher and less variable validation performance metrics across all models were kept as final test sets (see Methods). We did not include scMusketeers in this step to make sure that the test sets were selected independently from scMusketeers performances on them.

We selected highhly variable genes (HVG) as input to the models rather than using whole transcriptome data. Previous benchmarks suggest that the use of HVG performs in general evenly or better for the tasks of integration and automatic annotation, including for Deep Learning models such as scANVI [1, 2]. 3000 highly variable genes were selected using the method described by Sikkema & al [24], which favors genes that vary in every batch of a dataset (see Methods). The only exception was for UCE since it is based on a pangenome pretrained model.

## Supplementary method

### 9.6 Detailed model architecture

We consider a dataset of size *c* cells × *g* genes with *n*_*ct*_ cell types and *n*_*b*_ batches. We note its count matrix as ***X***, its cell types as ***C*** and its batches as ***B***. The default architecture of scMusketeers takes a log-normalized count matrix as input which we note 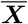

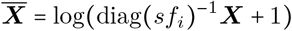

where diag (***sf***_***i***_)is the diagonal matrix composed of the size factor for each cell, which is equal to the library size divided by the median library size across cells. Library size is defined as the total number of counts per cell.

The scMusketeers model is an Autoencoder composed of three branches : a decoder, a cell type classifier and a batch declassifier in the form of a Domain-Adaptative Neural Network (DANN). The default configuration of the architecture is as follows :

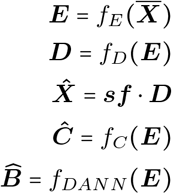

where *f*_*E*_, *f*_*D*_, *f*_*C*_, *f*_*DANN*_ are 2-layers MLP and ***sf*** is the vector of size factor computed on the input cells. The first layer of *f*_*DANN*_ is a gradient reversal layer, which inverts the sign of the gradient during backpropagation, making it a declassifier. ***sf*** can also be set to a unit vector to not explicitly model the size factor effect. ***E*** is the encoded layer of size *l* which consitutes the latent space. Then, ***D*** is the decoder’s output, 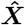 is the reconstructed count matrix, ***Ĉ*** is the classifier’s output and ***B*** is the DANN’s output.

By default, all layers use relu activation. Exceptions are made for the last layer of *f*_*E*_ which gives the encoded space and uses a linear activation by default, as well as the last layer of *f*_*D*_ which uses a relu to better capture the non-negative nature of the count matrix. A relu can be used instead for *f*_*E*_ to get a non-negative sparse latent space. Batch normalization and dropout are used by default for every layers of the model.

### 9.7 Permutations and training schemes

From this architecture, we deduce two losses from both classifiers. We using the categorical cross-entropy loss (***CE***) function for the batch removal declassifier. For the cell type classifier, we use the focal cross-entropy loss (***FCE***) to better account for the class imbalance inherent to single-cell RNAseq datasets.

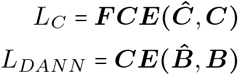

We used the Keras implementation of the focal categorical cross-entropy loss which is based on the work of [19]. Let us consider 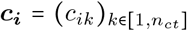, the one-hot encodedthe one-hot encoded true cell type vector of cell i and 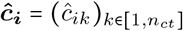, the predicted cell type probability output of the model where *c*ĉ_*ik*_ represents the probability that cell *i* belongs to cell type *k*. The focal loss is defined as :

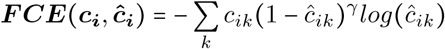

Where 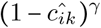 is the moduling factor, which gives a bigger weight to the poorly detected classes. *γ* is a focusing parameter which calibrates the strenght of this factor. It is set to 2 in the default configuration of scMusketeer.

For the reconstruction loss, we developed the concept of permutated reconstruction. Rather than comparing the reconstructed count of a cell to its original count, we compare it to a random other cell which belongs to the same cell type. Let 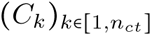 be the set of cells belonging to each cell type *k* and 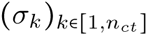 be a set of random permutations of *C*_*k*_ respectively. We then define *σ* = *σ*_*1*_ ○ *σ*_*2*_ ○ … ○ *σ*_*k*_ such that *σ*(*X*) = *σ* (⊍ *C*_*k*_) = ⊍ *σ* (*C*_*k*_). At each epoch, we draw a different set of *σ*_*k*_ and use it as the reference value to compare with the reconstructed value. We then use a mean squared error (***MSE***) loss :

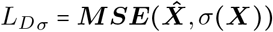

In some cases, we might also want to use a non-permutated reconstruction loss :

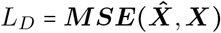

In all the training schemes further described, optimization is performed using gradient descent with the adam optimizer. While the classifier branch can only function in a supervised setup, the DANN and the reconstruction branch can function in a semi-supervised or unsupervised setup. We can propose several different training modes for our model by combining the different losses.

We define a “warmup” training mode to smooth out the batch effect. To do so, we consider the DANN loss and the reconstruction loss with permutation. Here, the total loss is :

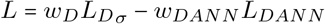

The whole dataset is passed through the model during training in a semi-supervised manner. To allow permutation for the reconstruction loss, we define pseudolabels for the unnanotated cells. If the model is not yet trained, we predict those pseudolabels by training a SVM on the annotated cells in the PCA space. When the model has been trained once, its predictions can be used as pseudolabels in a second round of training for further tuning. Another possible configuration is to use the permutation on annotated cells and to reconstruct unnanotated cells as themselves, in a standard autoencoder fashion. The negative contribution of the DANN loss to the total loss contributes to an alignement of the batches. Because the permutations are made across all batches, they additionally contribute to batch mixing. The intra-cell type permutations also encourages cells from similar cell types to group close-by in the latent space.

The main training mode of scMusketeers uses the full model in a supervised manner with a weighted combination of every losses. The loss is then defined as :

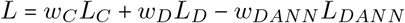

which is optimized in an adverserial manner to minimize the classification and reconstruction loss as well as maximize the batch effect. In this case, we don’t use permutation on the reconstruction loss, as it seemed to make the training procedure less stable.

To further refine the classifier, we can freeze all weights of the model apart from the classifier branch and train the classifier in a supervised manner on the latent space. The loss here consists of only the classifier loss. The loss weights are optimized during hyperparameters search.

We tested several different training schemes against one another, combining those training steps in different orders, with or without permutations and kept the top performing as default. We first warmup the model for a fixed number of epochs with the warmup training mode in an semi-supervised way with pseudolabels. We then switch to the full model training mode which we stop based on an early stopping on the validation balanced accuracy. We then train the classifier branch on the latent space, stopping based on an early stopping on the validation balanced accuracy. A second round of training can be performed using the same procedure, but using the predictions of the trained model as pseudo-labels for the warmup.

### 9.8 Selection of train and test dataset

We need to split each dataset in a train and a test split in proportions 80/20. Since we’re interested in training on different batches, this split is performed by taking out 20% of the entire batches as test and keeping 80% of the batches as our training set. Note that this may not result in a 80/20 split in terms of number of cells, as some batches are larger than others.

We then performed the benchmark of all methods but scMusketeers to determine the most stable test dataset. To do so, we split the remaining training batches in training and validation sets in proportions 80/20. We repeat the process 5 times using the sklearn cross validation function GroupShuffleSplit which can yield overlapping splits. The test and the validation data are treated as unnanotated data by the models which means that supervised method don’t use them at all during training whereas models with a semi-supervised component used them as unlabelled data.

## Appendix A Extended Data

**Fig. A1.**
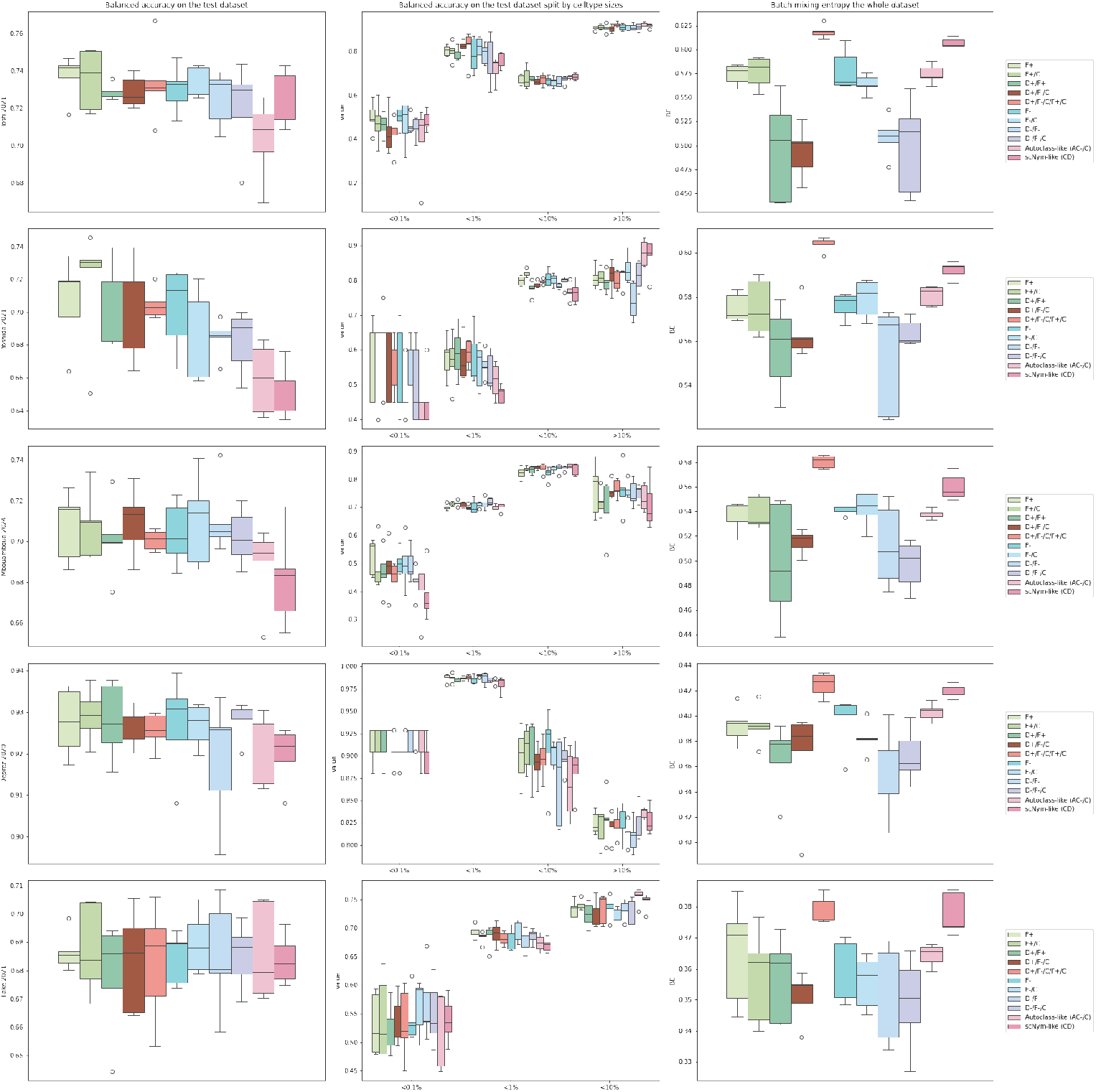
Performances for various training schemes tested on the Tosti, Yoshida, Mbouamboua, Deprez and Lake dataset. The schemes in legend correspond to the nomenclature described in Figure 1B

**Supplementary Table 1.**
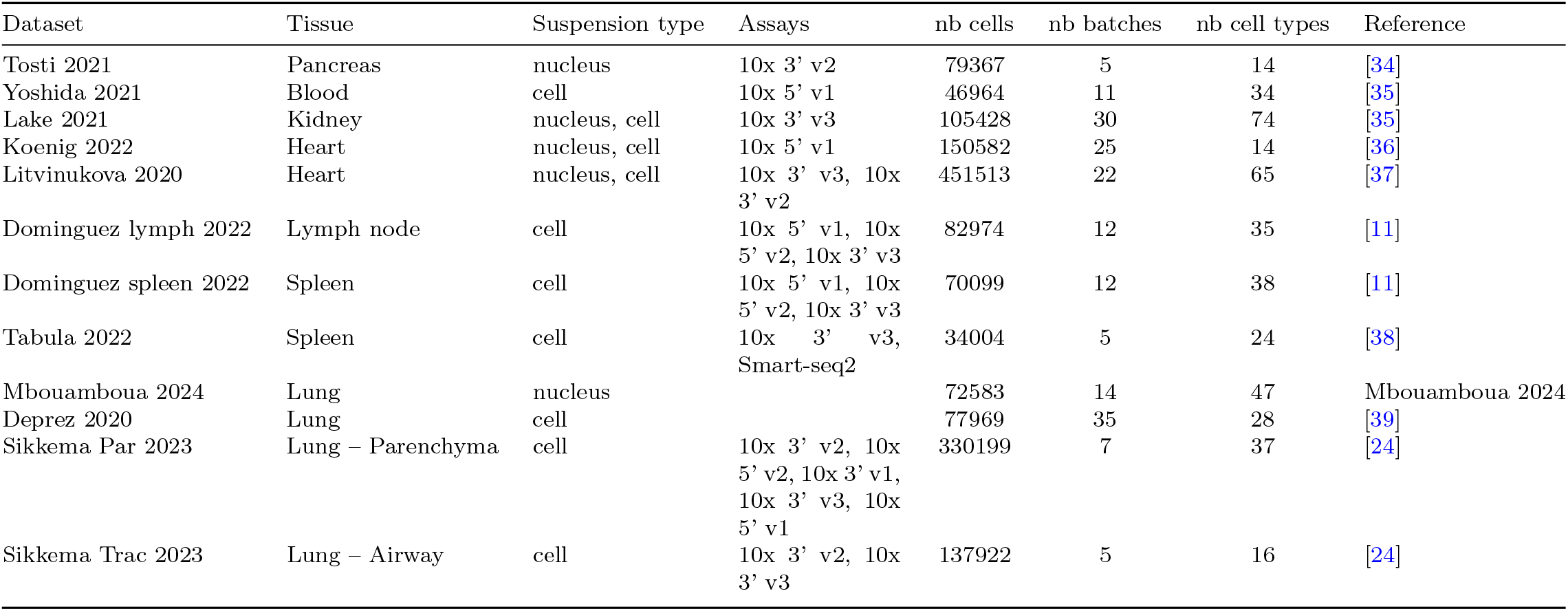
Datasets used for benchmark.

**Supplementary Table 2.** Models used for benchmark.

| Method | Type of model | Annotation | Embedding | Reference |
| --- | --- | --- | --- | --- |
| scANVI | Variational Autoencoder | Yes | Yes | [4, 18] |
| UCE+SVM | Pretrained Transformer | No | Yes | [6] |
| Harmony+SVM | cell type-informed PCA | No | Yes | [5] |
| PCA+SVM | PCA | No | Yes | [2] |
| Celltypist | Logistic Regression | Yes | No | [11] |
| scmap cells | Neirest Neighbors | Yes | No | [10] |
| scmap cluster | Neirest Neighbors | Yes | No | [10] |

**Fig. A2.**
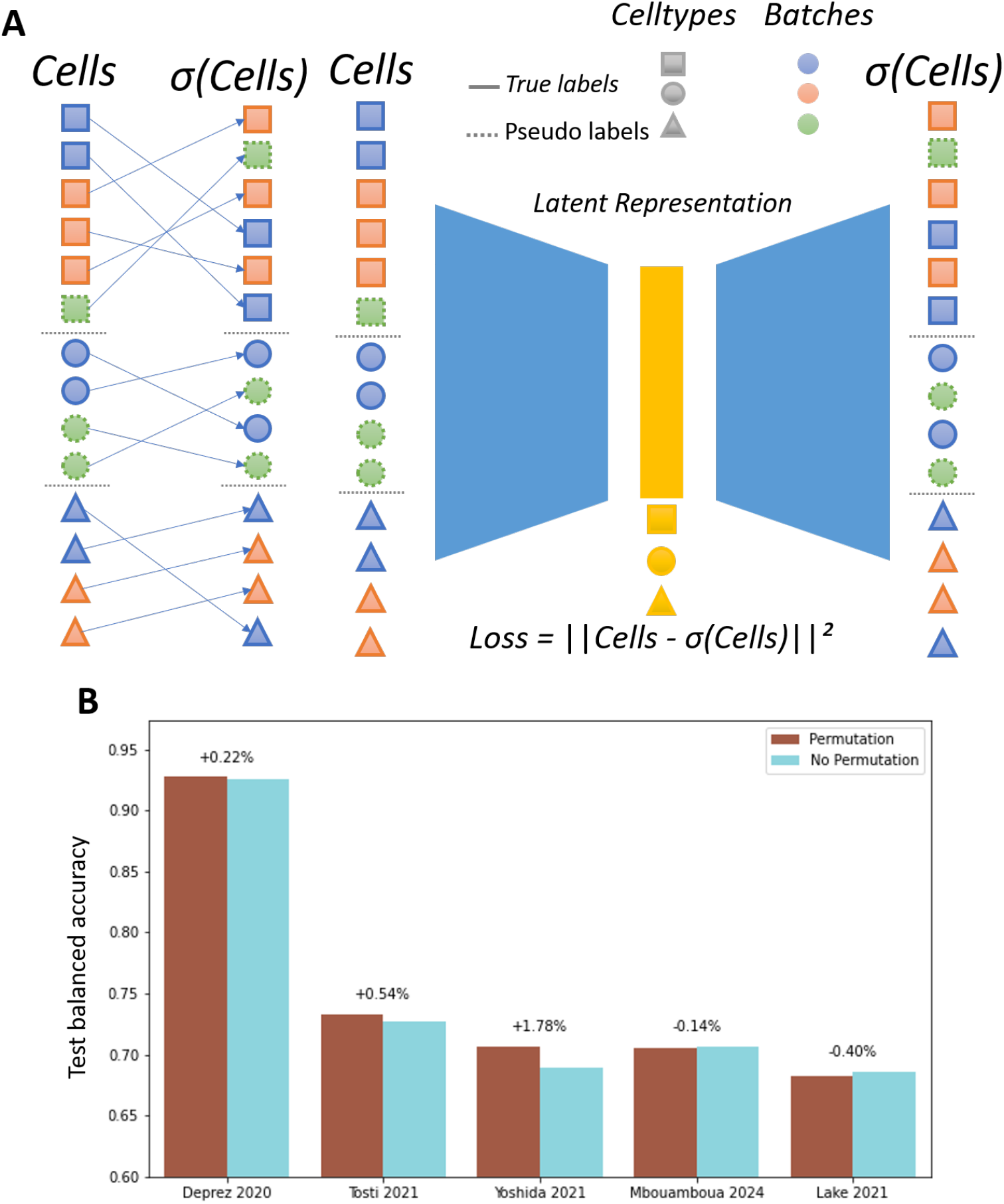
**A)** Illustration of permutated reconstruction. Each cell is matched with a different cell from the same cell type when computing the reconstruction loss. Permutations are made across batches, using pseudolabels for unannotated cells. The latent space is therefore expected to capture a batch-free representation of cell type identity. **B)** Barplots showing the mean balanced accuracy depending on the use of permutations. Mean accuracies were computed across four training schemes, F+, F+/C, D+/F+, D+/F-/C and their non-permutation counterparts.

**Fig. A3.**
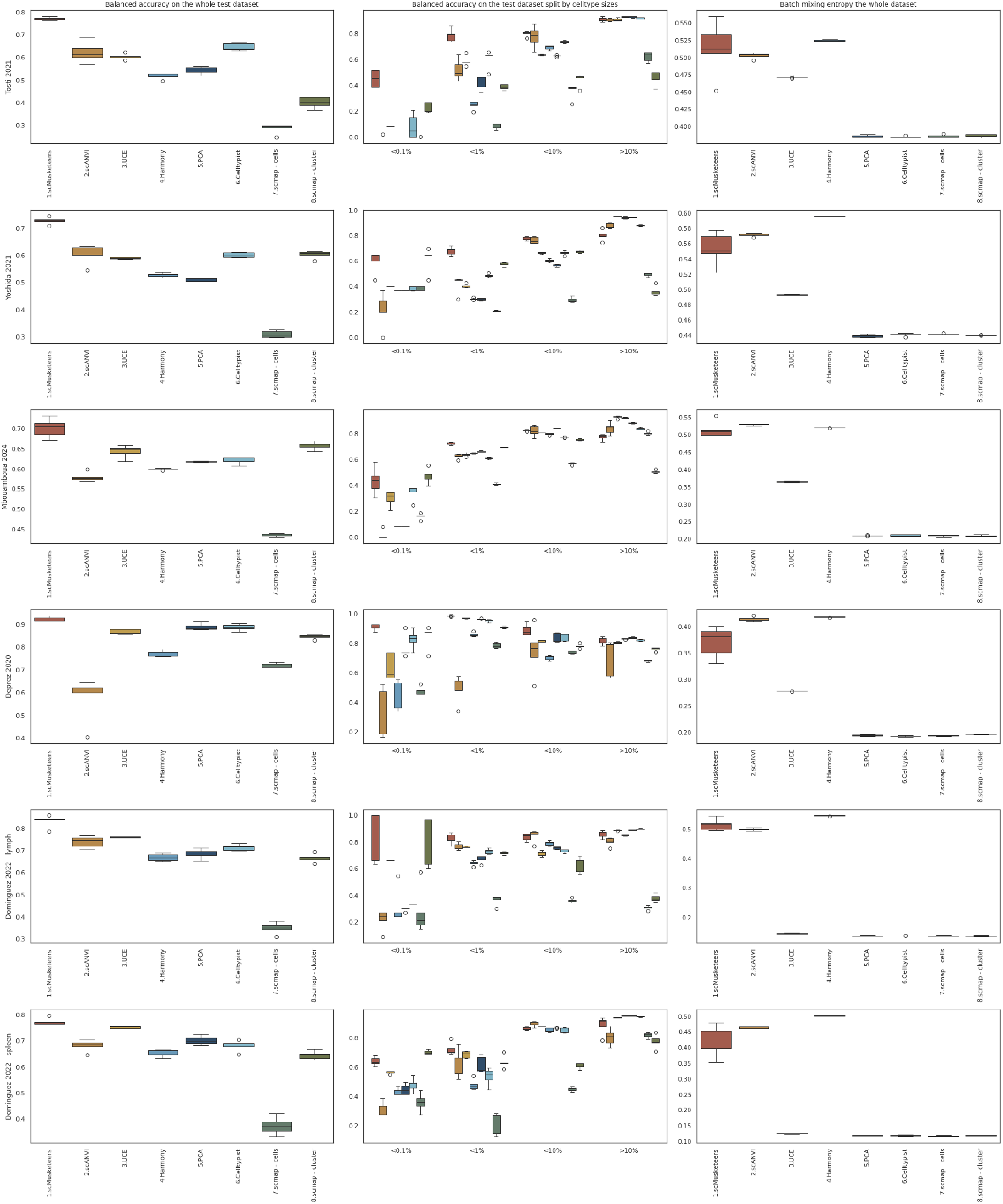
**A)** Test balanced accuracy (left), test balanced accuracy by cell type size (middle) and batch mixing entropy on the entire dataset (right) when training on a subset of the batches and predicting on the rest. Each row is a different dataset. Datasets represented : Tosti, Yoshida, Mbouamboua, Deprez, Dominguez-Lymph node and Dominguez-spleen.

**Fig. A4.**
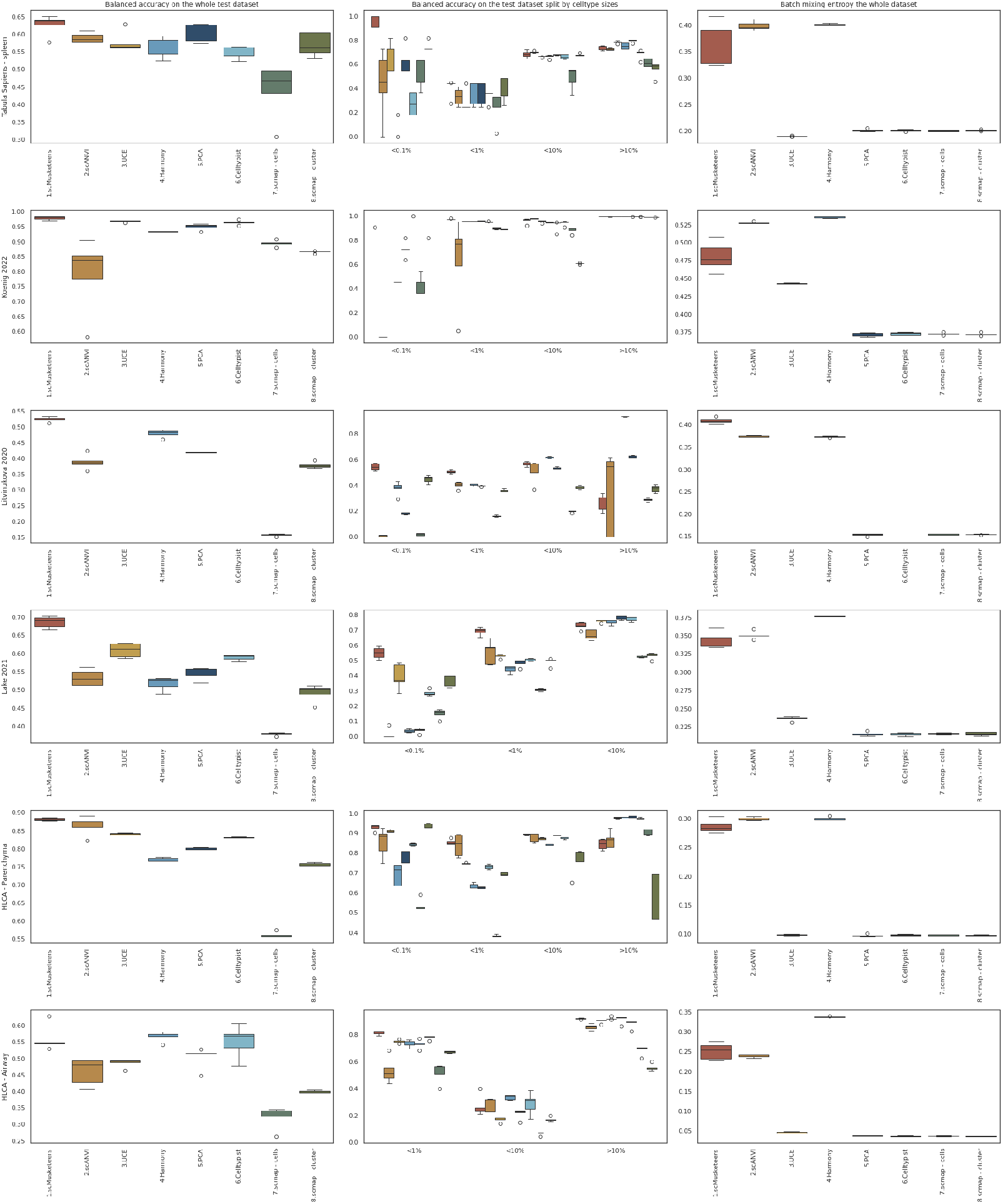
**A)** Test balanced accuracy (left), test balanced accuracy by cell type size (middle) and batch mixing entropy on the entire dataset (right) when training on a subset of the batches. Each row is a different dataset. Datasets represented : Tabula Sapiens, Koenig, Litvinukova, Lake, HLCA-Parenchyma and HLCA-Airway.

**Fig. A5.**
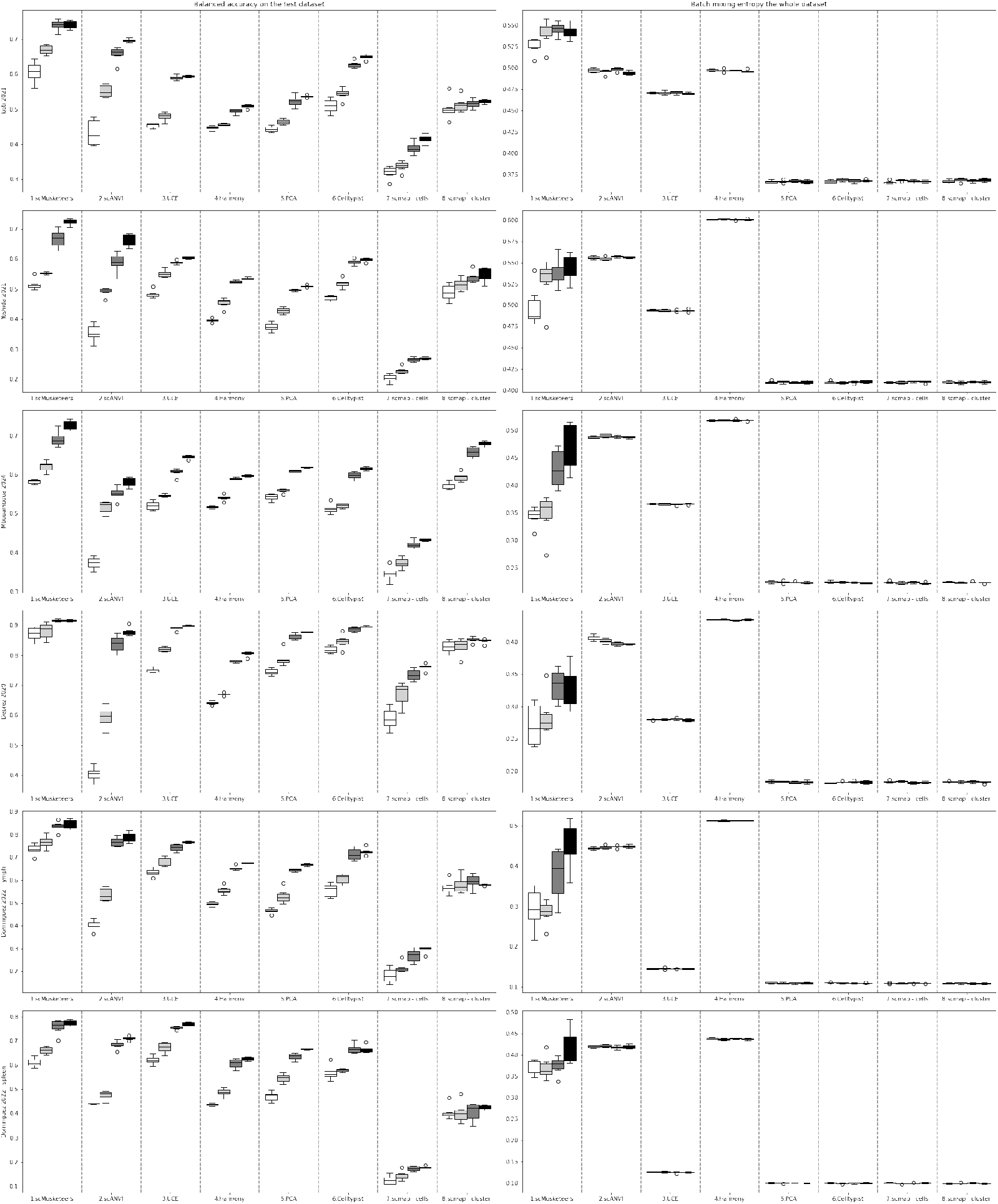
**A)** Test balanced accuracy (left), and batch mixing entropy on the entire dataset (right) when training on a percentage of all the cells. X-axis represents the percentage of the dataset used for training. Each row is a different dataset. Datasets represented : Tosti, Yoshida, Mbouamboua, Deprez, Dominguez-Lymph node.

**Fig. A6.**
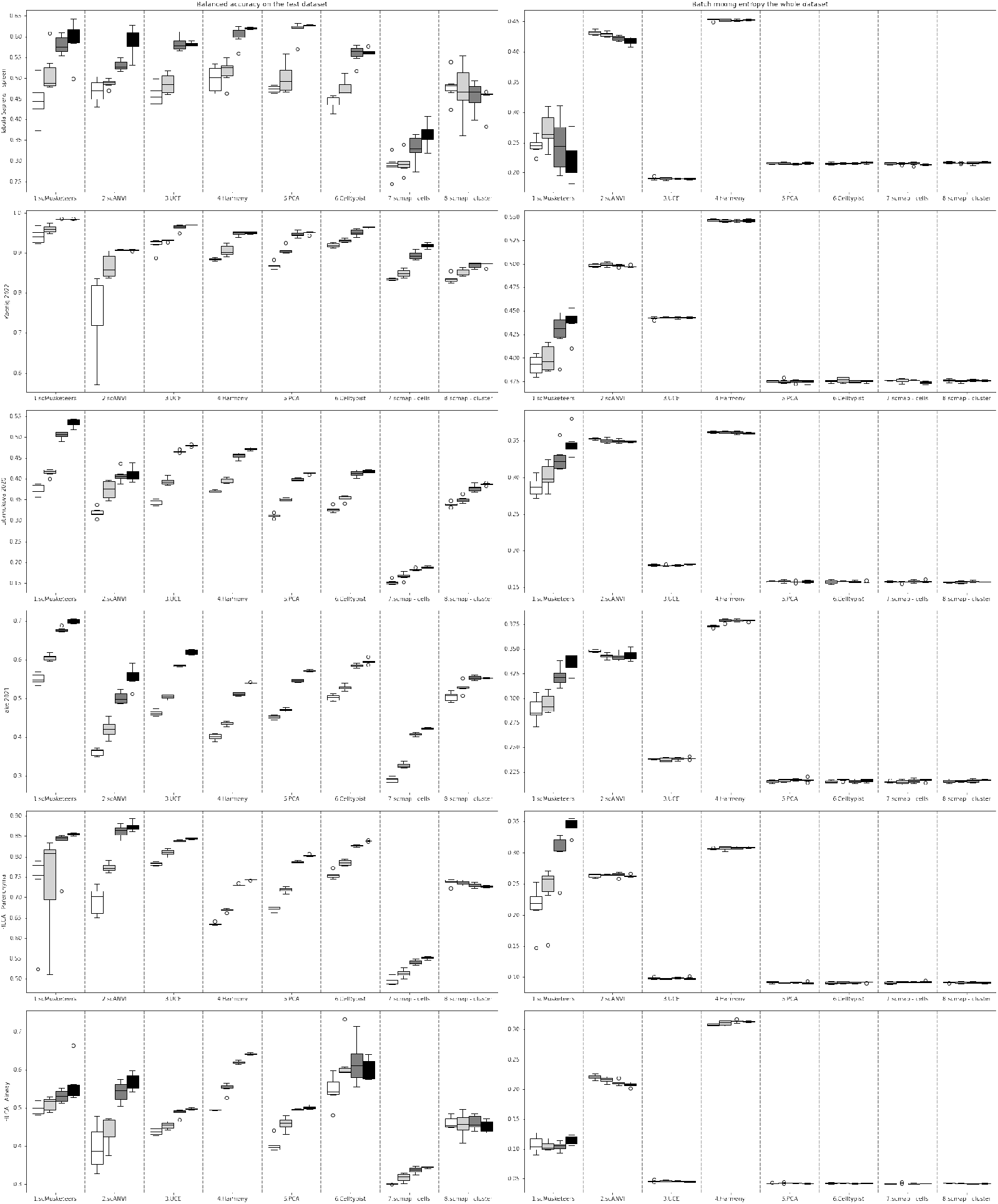
**A)** Test balanced accuracy (left), and batch mixing entropy on the entire dataset (right) when training on a percentage of all the cells. X-axis represents the percentage of the dataset used for training. Each row is a different dataset. Datasets represented : Dominguez-spleen, Tabula Sapiens, Koenig, Lake, HLCA-Parenchyma and HLCA-Airway.

**Fig. A7.**
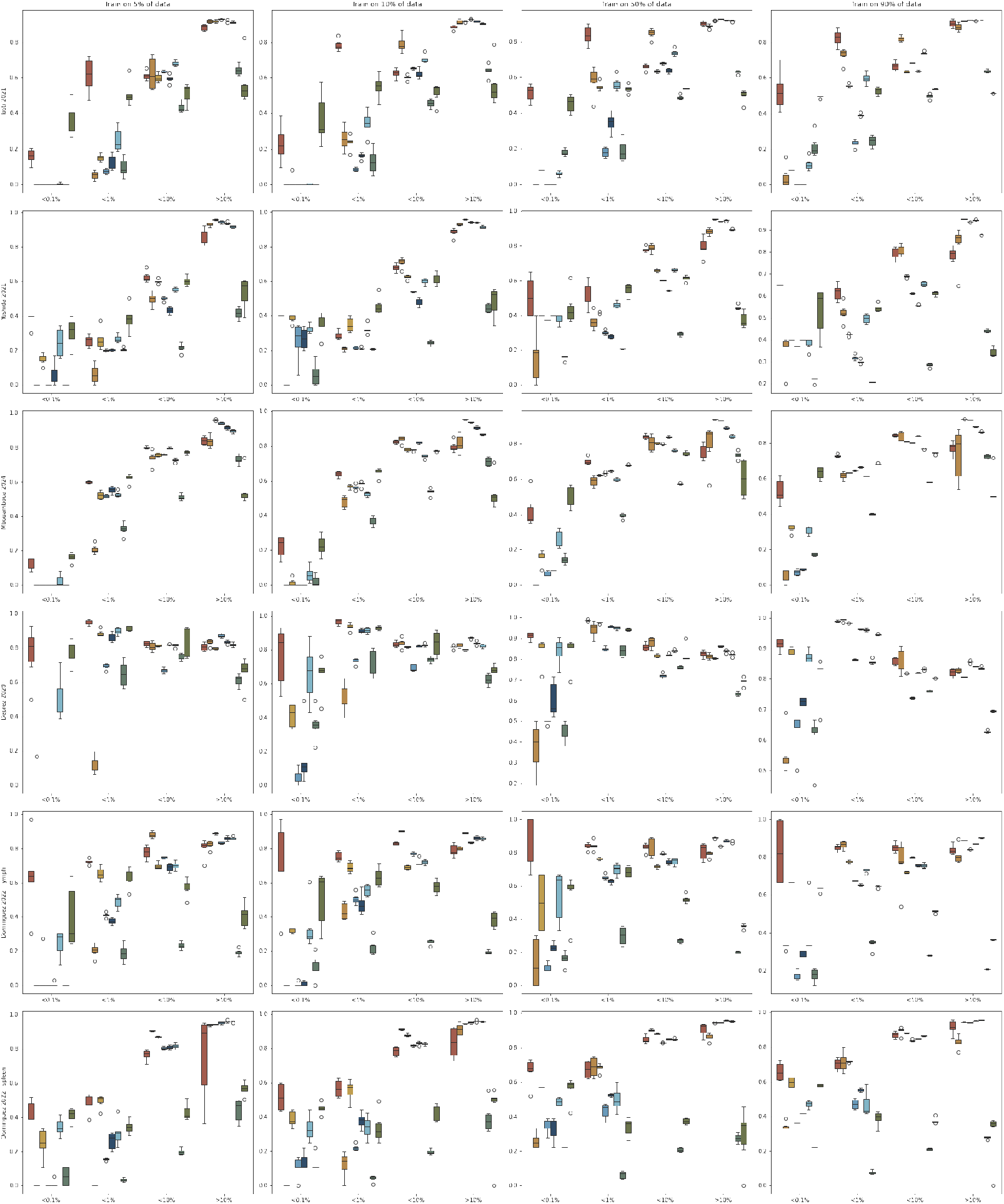
**A)** Test balanced accuracy depending on cell type size when training on a percentage of all the cells. Each column represents a different percentage of data used for training : 5%, 10%, 50% and 90%. Each row is a different dataset. Datasets represented : Tosti, Yoshida, Mbouamboua, Deprez, Dominguez-Lymph node.

**Fig. A8.**
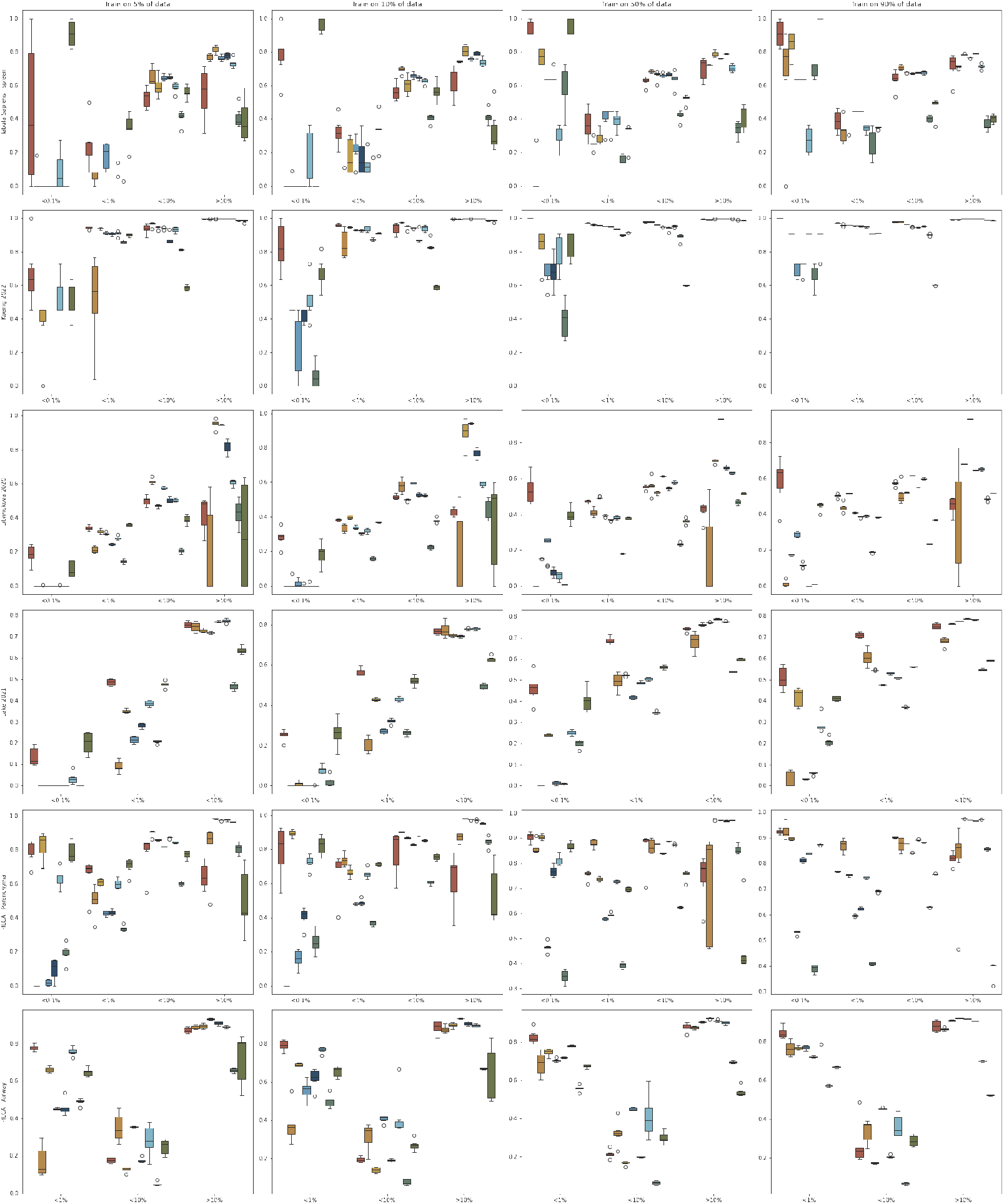
**A)** Test balanced accuracy depending on cell type size when training on a percentage of all the cells. Each column represents a different percentage of data used for training : 5%, 10%, 50% and 90%. Each row is a different dataset. Datasets represented : Dominguez-spleen, Tabula Sapiens, Koenig, Lake, HLCA-Parenchyma and HLCA-Airway.

**Fig. A9.**
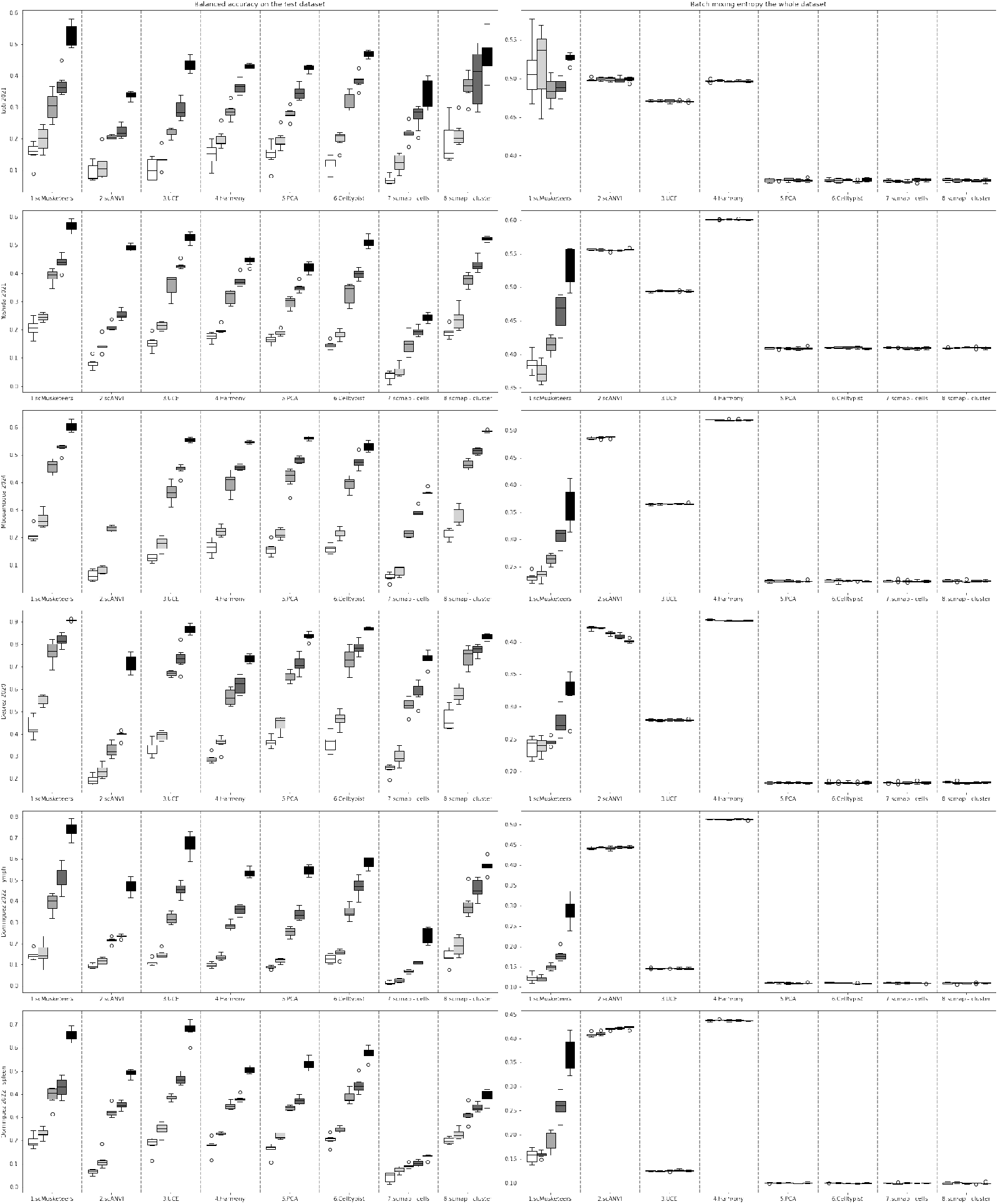
**A)** Test balanced accuracy (left), and batch mixing entropy on the entire dataset (right) when training on a fixed number of cells per cell type. X-axis represents the number of cells/cell type used for training. Each row is a different dataset. Datasets represented : Tosti, Yoshida, Mbouamboua, Deprez, Dominguez-Lymph node.

**Fig. A10.**
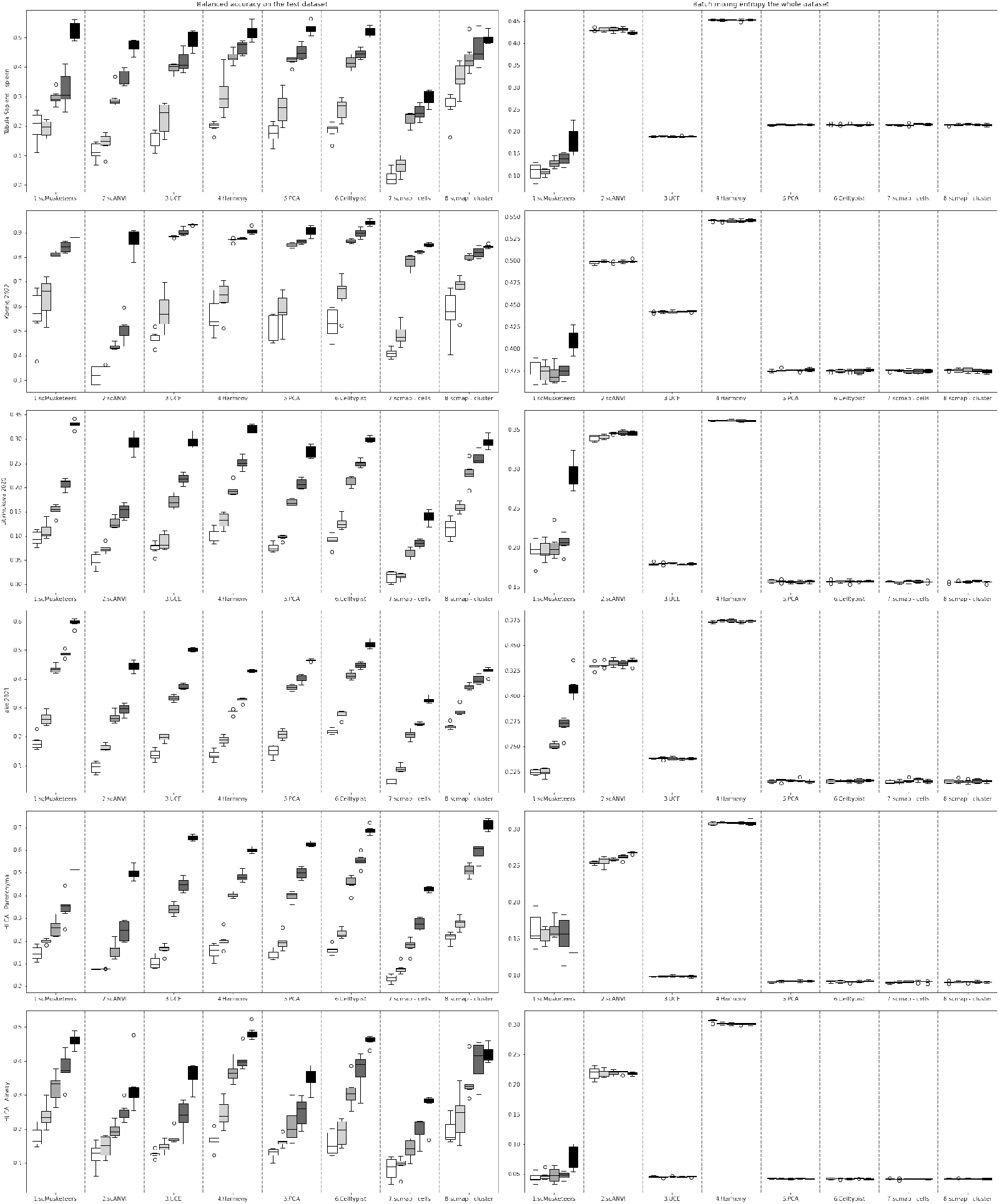
**A)** Test balanced accuracy (left), and batch mixing entropy on the entire dataset (right) when training on a fixed number of cells per cell type. X-axis represents the number of cells/cell type used for training. Each row is a different dataset. Datasets represented : Dominguez-spleen, Tabula Sapiens, Koenig, Lake, HLCA-Parenchyma and HLCA-Airway.

**Fig. A11.**
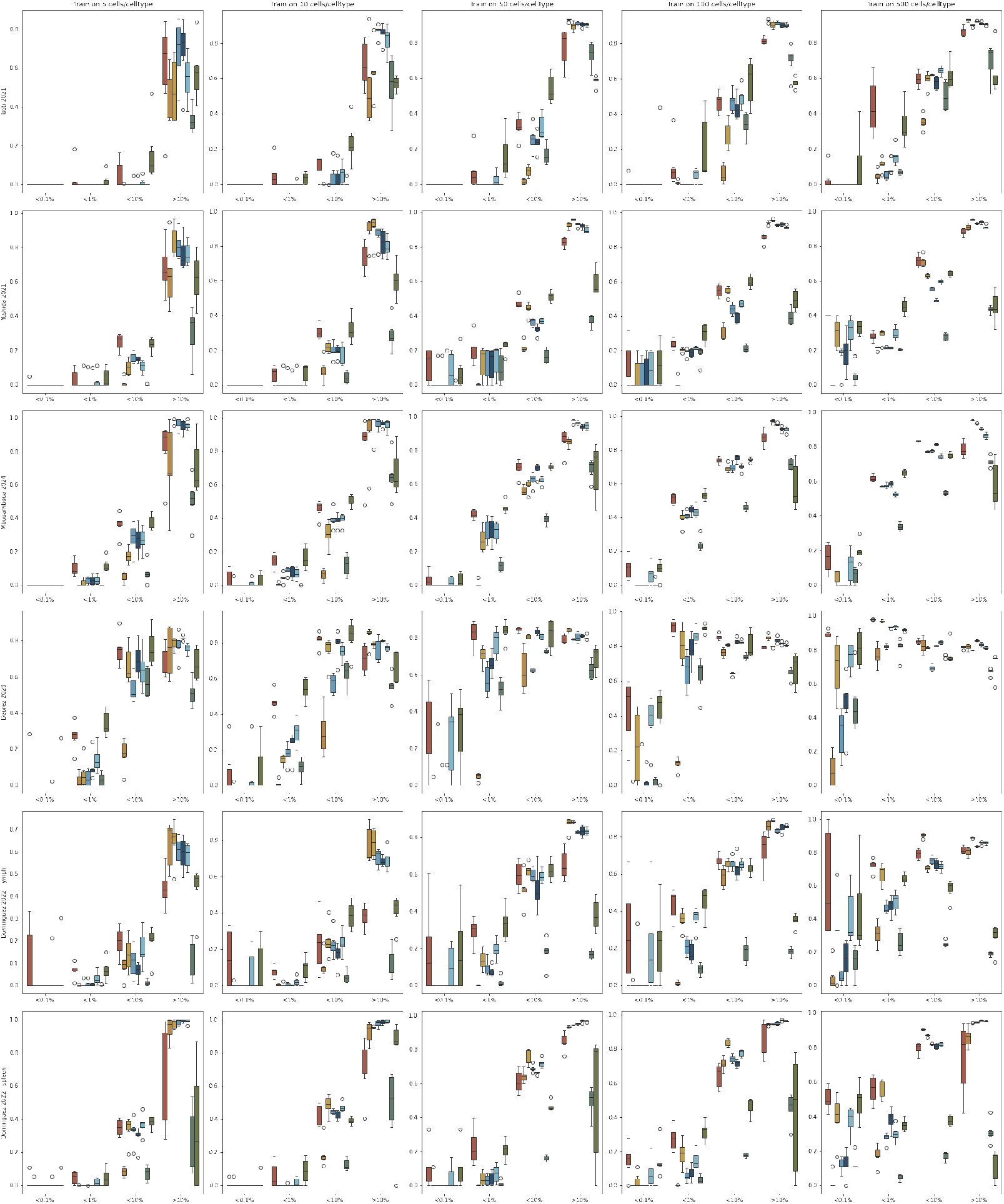
**A)** Test balanced accuracy depending on cell type size when training on a fixed number of cells per cell type. Each column represents a different number of cells per cell type used for training : 5, 10, 50, 100 and 500. Each row is a different dataset. Datasets represented : Tosti, Yoshida, Mbouamboua, Deprez, Dominguez-Lymph node.

**Fig. A12.**
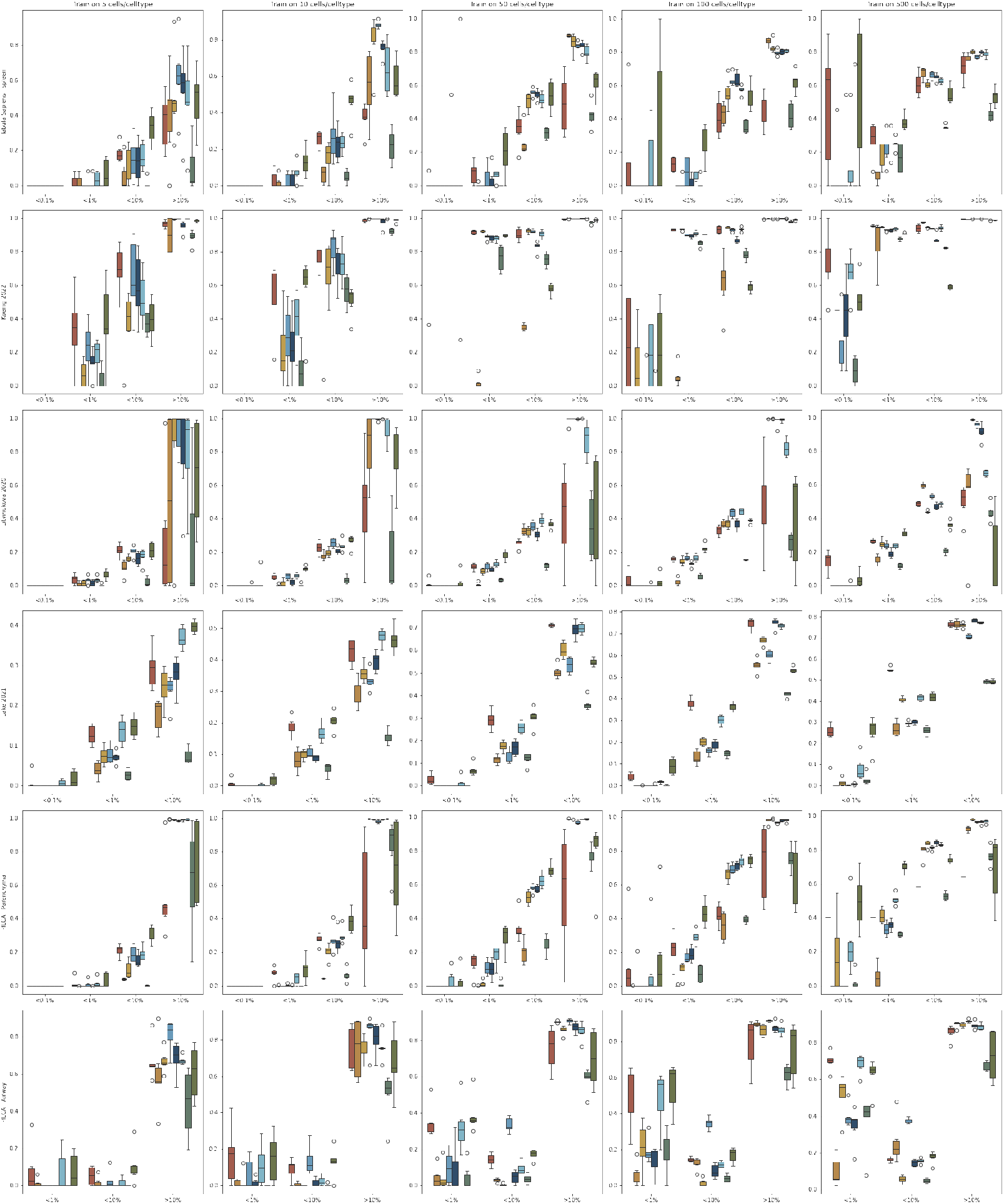
**A)** Test balanced accuracy depending on cell type size when training on a fixed number of cells per cell type. Each column represents a different number of cells per cell type used for training : 5, 10, 50, 100 and 500. Each row is a different dataset. Datasets represented : Dominguez-spleen, Tabula Sapiens, Koenig, Lake, HLCA-Parenchyma and HLCA-Airway.

**Fig. A13.**
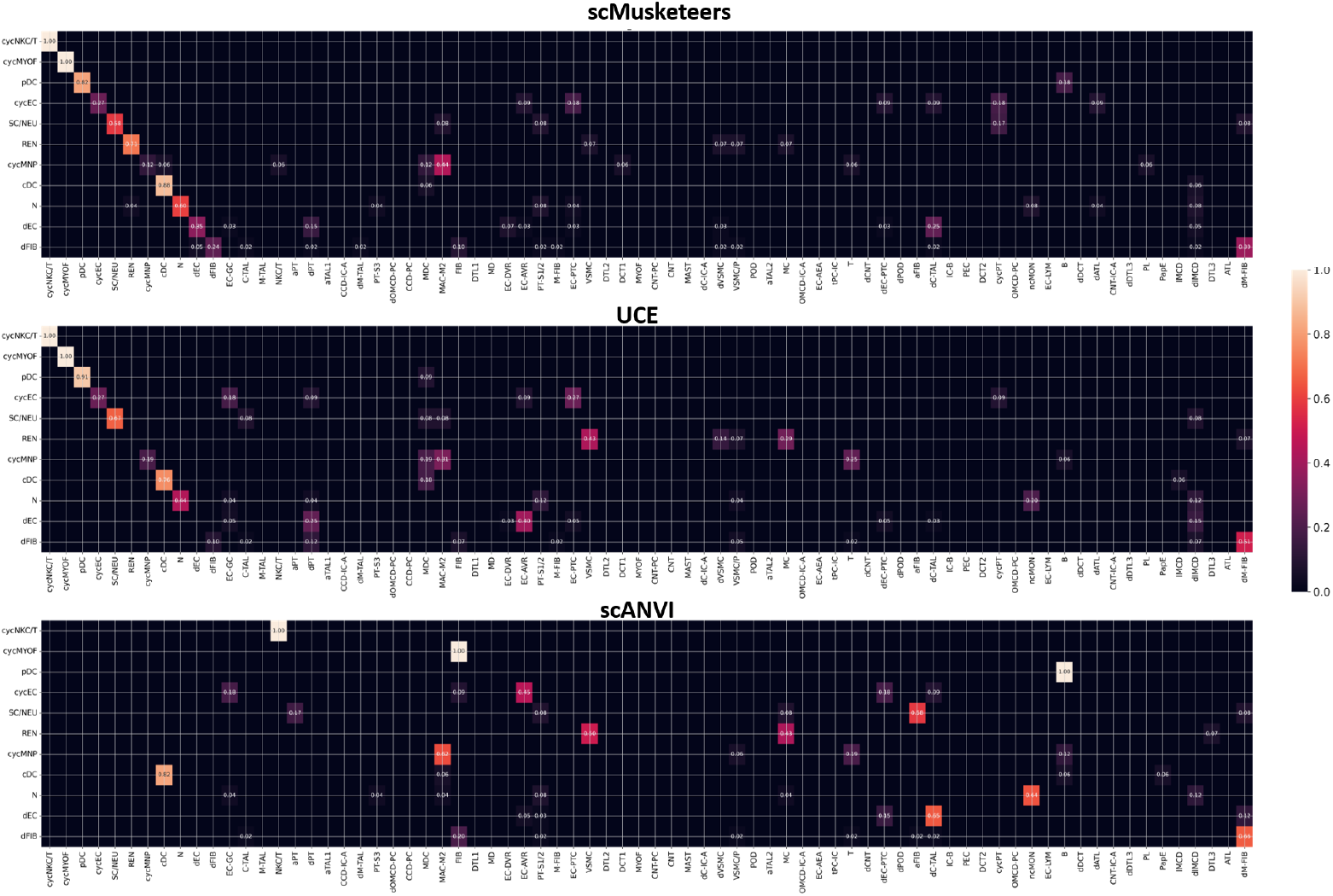
Cropped confusion matrices representing predictions of tiny cell types for one label transfer on the lake dataset.

**Fig. A14.**
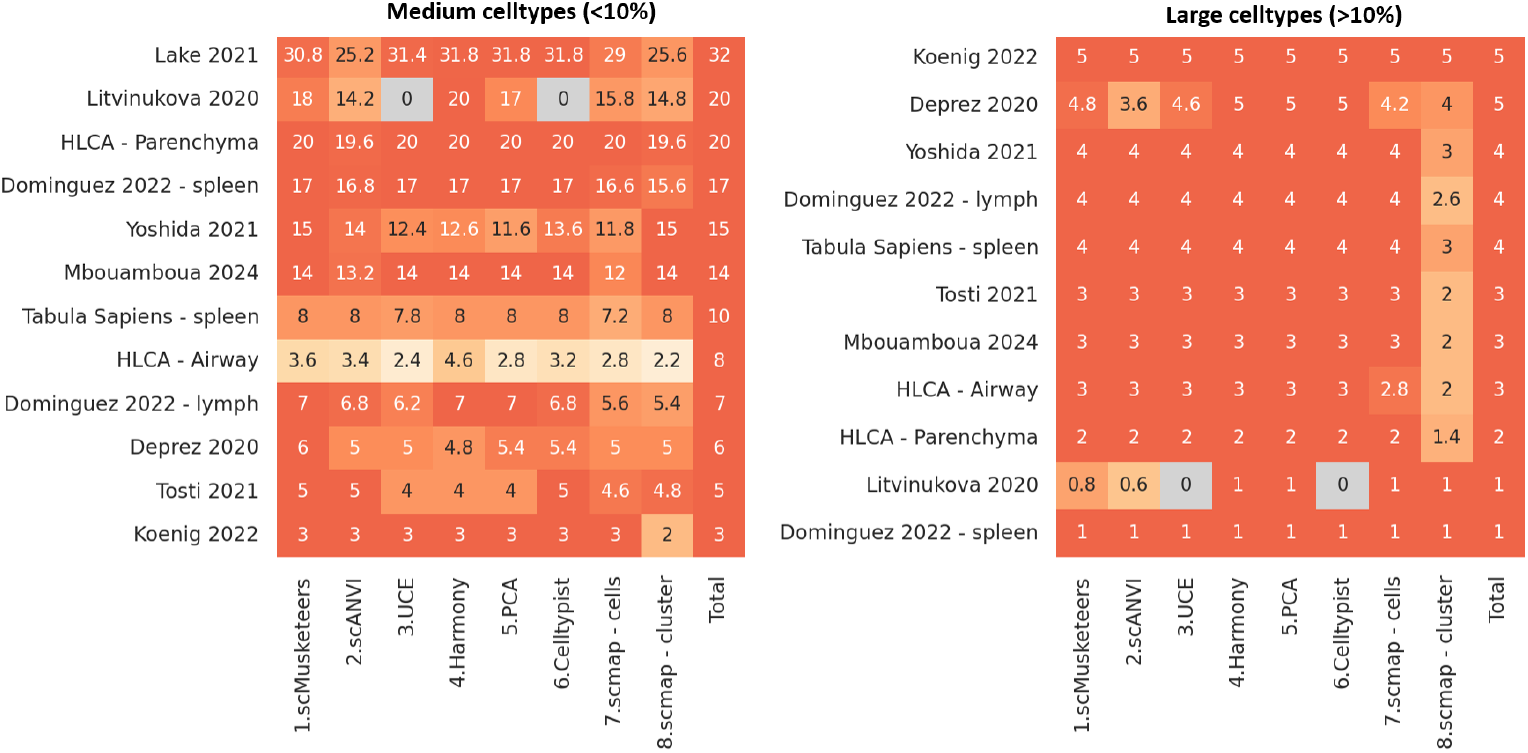
Detected cell types for large and medium size cell types. A cell type is detected if its accuracy is better than 20%

